# Constitutive differences in immune gene expression and energy storage phenotypes co-vary with winter environment in wood frogs

**DOI:** 10.1101/2024.12.11.627228

**Authors:** Grace J. Vaziri, Noah M. Reid, Tracy A.G. Rittenhouse, Daniel I. Bolnick

## Abstract

Many terrestrial ectotherms have gone to great evolutionary lengths to adapt to long cold winters; some have even evolved the ability to tolerate the freezing of most of the water in the body. Now, however, high-elevation, and high-latitude winters are experiencing an accelerated period of warming. Specialized winter adaptations that promoted fitness in a seasonally frozen environment may soon be superfluous or even maladaptive. We ask whether winter adaptations include changes in immune functions, and whether changing winter conditions could exert disparate effects on populations of a wide-ranging terrestrial ectotherm, the wood frog (*Lithobates sylvaticus*). By rearing wood frogs from ancestral winter environments that vary in length and temperature in a common garden, and reciprocally exposing post-metamorphic frogs to unfrozen and frozen artificial winter conditions in the lab, we were able to decompose transcriptomic differences in ventral skin gene expression into those that were environmentally induced (responsive to temperature), genetically determined, and those that varied as an interaction between genotype and environment. We found that frogs from harsh ancestral winter environments constitutively upregulated immune processes, including cellular immunity, inflammatory processes, and adaptive immune processes, as compared to frogs from mild ancestral winter environments. Further, we saw that expression of several genes varied in an interaction between genotype and artificial winter. We suggest that just as winter climates likely served as the selective force resulting in remarkable winter adaptations such as freeze tolerance, they also induced constitutive changes in immune gene expression.

## Introduction

Climate change is rapidly affecting winter conditions, which will exert new pressures on cold-adapted organisms (O’Connor & Rittenhouse, 2016; Williams et al., 2015; Zhu et al., 2019). Ectotherms, including amphibians, can be particularly sensitive to winter conditions. While behavioral thermoregulation in the warm months allows ectotherms to remain close to their thermal optima, behavioral thermoregulation is less effective during the cold months when environmental temperature heterogeneity is reduced (Muñoz & Bodensteiner, 2019) and when low temperatures hinder the mobility of ectotherms. To deal with winter conditions at high latitudes and altitudes, ectotherms must have adaptations to circumvent cold injury, infection, and starvation—hazards borne from cold temperatures, reduced physiological rates, and resource limitations. Such winter adaptations can be highly specialized, even allowing some vertebrates to tolerate prolonged and extensive freezing (Storey, 1992).

Research on the effects of changing winters on amphibians have addressed the direct influence of winter conditions on metabolism, energy usage, and survivorship (M. J. Fitzpatrick et al., 2019, 2020; O’Connor & Rittenhouse, 2016; Sinclair et al., 2013). For example, experimental evidence has shown that in the wood frog (*Lithobates sylvaticus*), overwintering survival is enhanced under conditions with consistent snow cover (O’Connor & Rittenhouse, 2016). Across the wood frog range, winter conditions vary in ways beyond snow cover, including winter length, soil temperature, and number of freeze-thaw cycles. This results in a mosaic of winter energy requirements. Local winter climates likely drive selection for energetic and physiological investment in winter adaptations among species that occupy a spectrum of winter environments (Costanzo et al., 2015). For example, in environments in which winters have historically been long and very cold, amphibians may accumulate energy and cryoprotectant reserves at the expense of somatic growth (amphibians do not follow Bergmann’s rule; Adams and Church, 2008). Consistent with this expectation, in the eastern US, wood frogs from northern populations tend to be (but are not always) smaller than those from warmer southern populations (Davenport & Hossack, 2016; Martof & Humphries, 1959).

To this point, wood frogs from Alaska have been shown to store proportionally more glycogen compared to Ohioan wood frogs in preparation for winter, indicating that these populations may be locally adapted to different winter conditions (do Amaral et al., 2013). Glycogen accumulation is a key adaptation, enhancing freeze tolerance and allowing wood frogs to occupy higher latitudes than any other North American amphibian. Wood frogs store glycogen in the liver and urea in blood plasma to increase their cryoprotective osmolytes during freezing (Costanzo et al., 2013). In wood frogs, extensive work has been conducted to demonstrate links between glycogen stores and the extent and duration of freeze tolerance (Costanzo et al., 2013; do Amaral et al., 2013, 2016). To date, however, few studies have explored the extent to which energy storage phenotypes are evolved or plastic, or whether they co-vary with physiological traits other than freeze tolerance. Answering these fundamental questions is essential for understanding the extent to which populations across species distributions will be affected by changing winter conditions as climate change progresses.

In addition to energy storage phenotypes, immune processes may also co-vary with winter climates. Understanding the evolution and ecology of immunity in the cold months is an important component of forecasting how climate change will influence terrestrial ectotherms (Ferguson et al., 2018). Ectotherms generally regulate physiological processes, such as immunity, according to a thermal performance curve (Cohen et al., 2017; Ferguson et al., 2018; Sinclair et al., 2016). When temperatures dip below an organism’s optimum, therefore, rates of immune processes among ectotherms are expected to decrease and indeed many, but not all, immune processes are downregulated in the cold (Cooper et al., 1992; Vaziri et al., 2024).

Recently, our group found that wood frogs reduce the expression of many genes in the skin and spleen that are associated with adaptive immune processes between early fall and midwinter, when temperatures are markedly colder (ref et al.). In contrast, Gupta and colleagues (2020) showed that several oxidative stress-response genes are upregulated in the liver and skeletal muscles of wood frogs in frozen conditions. Additionally, some components of the innate immune system have been shown to be maintained or upregulated in response to cold exposure (Cooper et al., 1992; Maniero & Carey, 1997), suggesting that ectothermic hosts do not totally forsake investment in immune defense in the winter.

The vertebrate immune system comprises a massive number of pathways, cell types, tissues, and even behaviors; these components may respond differently to thermal and other environmental cues. Studies of tissues and organs that are directly involved with overwintering survival including adipose tissue, liver, and skeletal muscle, have led to insights with potential importance for clinical applications (Lee et al., 1992; Saxton et al., 2022). Additionally, other researchers have shown that the antimicrobial activity of wood frog ventral skin, an innate immune function, is enhanced by short term (24h) bouts of freezing (Katzenback et al., 2014). Researching how the gene expression of the skin responds to winter will broaden our understanding of how tissues not directly related to the ability of an organism to survive overwintering operate in cold conditions. In particular, investigating how the skin, an important immune tissue and mucosal barrier in amphibians, and the first barrier between an organism and the environment (Le Sage et al., 2021; Varga et al., 2019; Woodhams et al., 2000), is regulated during winter, could provide insight into environmental stressors—past and present—and current tradeoffs that surviving harsh winters entail.

Environmental heterogeneity across the geographic range occupied by overwintering wood frogs may underlie the evolution of local energetic and immune adaptations. While frozen, overwintering frogs produce energy using anaerobic metabolism (Al-attar et al., 2020; Storey, 1987; Voituron et al., 2002); unfrozen overwintering frogs employ fatty acid oxidation (lipid metabolism)(L. Fitzpatrick, 1976; Long, 1987). Anaerobic (carbohydrate) metabolism, including glycolysis, enhances production of many inflammatory cytokines; conversely lipid metabolism can moderate the inflammatory functions of macrophages (O’Neill et al., 2016). Origin winter environments may drive variation in the proportion of ingested energy that frogs allocate towards glycogen versus lipid accumulation (Costanzo et al., 2013). We hypothesize that the metabolic pathways favored by frozen versus unfrozen wood frogs will promote different immune phenotypes, and as such, origin winter environments that have selected for different energy storage phenotypes may also underlie locally adapted immune gene expression profiles among populations. Energy allocation strategies necessitated by different winter conditions and different cellular metabolic pathways favored while frozen versus unfrozen may underlie different immune phenotypes based on both host genetic background and ambient winter conditions. We use an experiment to evaluate if wood frogs regulate their overwintering immunity, depending on both their origin winter environment (warmer and shorter or colder and longer), and as a short-term plastic response to the winter environment they experience. In doing so, we also investigate one potential mechanism of the differences in observed immune regulation, differences in energy storage phenotypes.

We used a common garden experiment to test whether adaptations to origin winter environments, or plastic responses to current winter temperatures, influence pre-winter energy allocation strategies and winter immune phenotypes. We reared frogs from locations where winter conditions differ in length and temperature but incur comparable energetic costs for wood frogs. Then, we reciprocally exposed frogs to artificial winter treatments in which temperatures were either above or below freezing (0°C; Figure 1B). We hypothesized that skin immune gene expression would vary based on winter conditions of their ancestral population origins, and that such a relationship would be mediated by whether expression is promoted by glycolysis or lipid metabolism, and with respect to an interaction between artificial winter treatment temperature and origin winter environments (‘harsh’ or ‘mild’). This hypothesized interaction is because the ability to produce different immune measures (and the benefit gained by doing so) depends on how stored energy is allocated—a physiological trait that may vary geographically—and the metabolic pathways available to immune cells, which vary with whether frogs are hibernating in a frozen or unfrozen state. Specifically, we reasoned that the fitness benefit gained from enhanced freeze tolerance should vary with the winter conditions frogs experience. We predicted that frogs from populations that have adapted to long cold (harsh) winters would allocate proportionally greater amounts of energy toward glycogen accumulation. Conversely, we predicted that those from populations that have historically experienced shorter, less cold (mild) winters would allocate proportionally more energy toward lipid accumulation than those from harsh origin winter environments.

**Figure 1.**
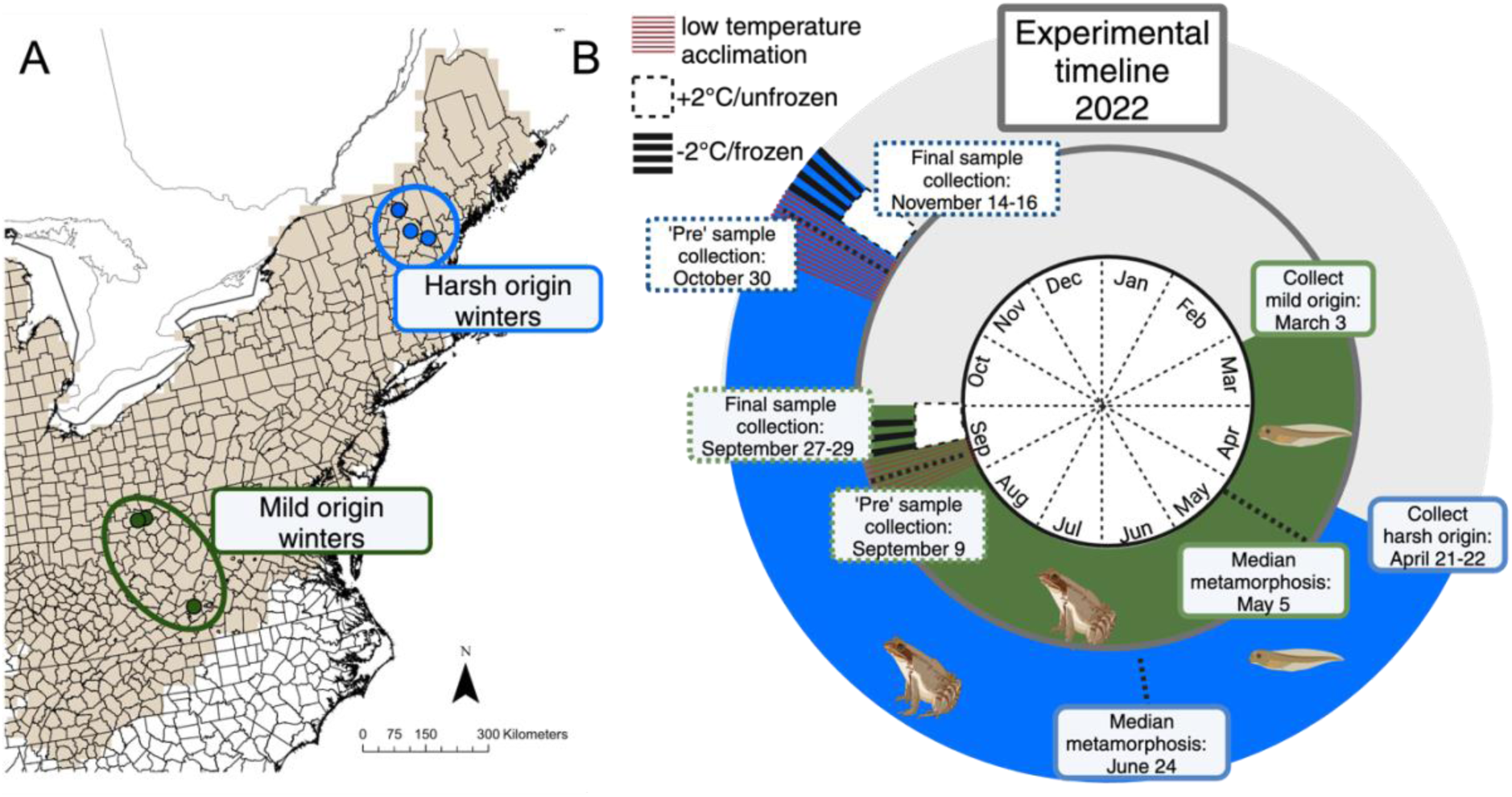
A) Wood frog range in the eastern United States (brown). Sampling sites are represented with colors; sampling sites had equivalent winter energy requirements, but different ancestral winter conditions. Harsh (long and cold) winters in blue, Mild (shorter warmer) winters in green. B) Experimental timeline: Egg masses were collected in March and April from locations places where wood frog populations have experienced shorter and warmer, or longer and colder winters, respectively. Frogs were reared in a common garden (laboratory conditions) for identical periods of time to obviate chronological mismatches in phenology between frog populations from northern and southern collection sites.

## Methods

| Scale of inference | Scale at which the factor of inference is applied | Number of replicates at the appropriate scale |
| --- | --- | --- |
| Winter origin environment<br>(site type) | population | 3 |

### Population selection

We collected egg masses from populations within the Eastern mtDNA clade (Lee-Yaw et al., 2008). Using climate data spanning 1980-1999 from the eastern extent of the wood frog range (M. J. Fitzpatrick et al., 2020) we performed a suitability analysis to identify 25-km^2^ areas from which to collect egg masses (Figure 1A). We used M.J. Fitzpatrick and colleagues’ (2020) definition of winter length—the period during which frogs are restricted to their hibernacula, as well as their simulated estimates of energy expenditure (in J) for a 11.6 g adult wood frog throughout the duration of a winter, and their estimates for the number of freeze thaw cycles, and temperature at 5-cm soil depth. Potential sites were identified using the following criteria: 1) Average energy required by a wood frog to survive winter at a site was within half a standard deviation of the mean energy required across all sites in the dataset (11,892±979 J). 2) Sites experienced between 7-11 freeze-thaw cycles per winter on average. 3) Sites had average winter soil temperatures (5-cm soil depth) of ≤-0.5°C, or ≥0.5°C. After filtering according to these criteria, we used a combination of iNaturalist observations of breeding wood frogs, Google Earth searches, and word-of-mouth to identify sites from which to collect egg masses (Table 1). We performed Welch’s two-sample t-tests to confirm that the selected sites met our criteria of having similar energy requirements for winter survival (mean_mild_ =11,469 J, mean_harsh_=12,038 J, t = −1.7, p = 0.19), similar numbers of freeze-thaw cycles per winter (mean_mild_ = 7.2 cycles, mean_harsh_=8.7 cycles, t = −1.5, p = 0.3) and were primarily distinguished by whether winter conditions were harsh (blue-circled sites, Fig. 1; mean_winter length_= 170.5 days, mean_temperature_ = −1.1°C), or mild (green-circled sites, Fig. 1; mean_winter length_= 116.9 days, mean_temperature_ = 0.6°C, t_winter length_ = 10.2, p_winter length_=0.003, t_temperature_ = −46.5, p_temperature_ < 0.0001). Up to three partial (1/4 of total) egg masses were collected from three ponds in the mild winter areas, and from three ponds in the harsh winter areas (all ponds separated by at least 1-km; Table 1).

**Table 1.** Collection data for wood frog egg masses used in the study.

| Winter<br>cat. | Population<br>name | Latitude | Longitude | Collection<br>date | *Winter<br>energy use<br>(J/year) | *Winter<br>soil temp<br>(°C) | *Num.<br>freeze-thaw<br>cycles | *Snow<br>cover<br>(cm) | *Winter<br>length<br>(days) |
| --- | --- | --- | --- | --- | --- | --- | --- | --- | --- |
| Mild | Burke | 37.322600 | -80.372140 | 03/03/22 | 12031.54 | 0.63 | 10.64 | 2.33 | 115.7 |
|  | ORI | 39.403278 | -81.205888 | 03/03/22 | 12266.62 | 0.59 | 7.41 | 3.64 | 120.84 |
|  | Barr | 39.387920 | -81.423410 | 03/03/22 | 11816.37 | 0.63 | 8 | 2.94 | 114.12 |
| Harsh | Victory | 44.544630 | -71.851910 | 21/04/22 | 11485.04 | -1.12 | 7.09 | 25.63 | 174.36 |
|  | Woodstock | 44.031010 | -71.693470 | 21/04/22 | 11986.75 | -1.02 | 7.08 | 28.41 | 176.4 |
|  | Coombes | 43.765390 | -71.250550 | 22/04/22 | 10936.46 | -1.02 | 7.33 | 27.26 | 160.88 |

### Husbandry

We reared egg masses to Gosner stage 25 (free-swimming larvae) in environmental chambers initially set to the average temperature of the water bodies from which egg masses were collected, 5.5°C, and gradually warmed water to 20.5°C degrees (∼0.6°C/day). Egg masses were maintained in 5-L plastic containers filled with aged tap water treated with StressCoat (0.13mL/L; Cat #85, Mars Fishcare North America, Inc., Chalfont, PA, US). At Gosner stage 25 larvae were transferred to a laboratory maintained at 20.5°C, where they remained until experimental exposure to artificial winter conditions commenced. Larvae were stocked at two densities, 1 larva/L H_2_0, and 4 larvae/L H_2_0. The majority (74/92, 80%) of animals used in the final analysis of this study were reared at a density of 1 larva/L H_2_0, but we also used animals that were reared in the higher density environment to supplement sample sizes from each origin winter environment:artificial winter treatment combination. We performed all analyses (described below) on both the full group of samples, and on the subset that were reared in the lower density environment and found qualitatively equivalent results. Therefore, we present results from the full dataset in the manuscript.

We conducted full water changes every four days and fed larvae a 50:50 (by mass) blend of Tetramin Tropical flake (Cat #77101, Tetra, Blacksburg, VA, US) and Hikari algae pellets (Cat #21366, Hikari USA, Hayward, CA, US) ground to a homogenous powder. Larvae were fed daily to approach *ad libitum* feeding conditions without fouling the water. At Gosner stage 42 (emergence of one or both forelimbs), metamorphs were transferred from aquatic tanks to small sandwich containers placed on an incline with a small amount (c.a. 50mL) of water so that metamorphosing animals could select whether to be in water or on a dry surface. Throughout the period of metamorphosis, larvae in aquatic tanks were consolidated (within populations) to maintain the original stocking density of the tank. When metamorphs had fully resorbed their tails (Gosner stage 46), they were transferred from sandwich containers to terrestrial tanks. Terrestrial tanks were stocked at a density of 3 metamorphs/gallon and were supplied with a layer of paper towels moistened with aged tap water and a layer of leaf litter for cover. Metamorphs were fed a mix of fruit flies, *Drosophila melanogaster* and *D. hydei,* every other day throughout the experiment. Prior to one feeding each week, fruit flies were dusted with enough vitamin and mineral supplement powder to completely cover their bodies (Cat # 7-88286-00200-9, Rep-Cal Research Labs, Los Gatos, CA, US). Terrestrial tanks were cleaned every four days.

### Experimental winter

At 17 weeks post-metamorphosis, when all frogs were still sexually immature, a pre-winter sample was collected from each group (N = 17 mild origin, N = 14 harsh origin) to assess pre-winter energy storage (liver and frog carcass), and baseline gene expression (ventral skin; Figure 1B). Sample collection dates were asynchronous for frogs from different origin winter environments; frogs from mild origin winter environments were collected earlier in the year than those from harsh (see the offset in timing in Figure 1B). To deal with the asynchrony, all frogs were reared for the same number of weeks post-metamorphosis. Remaining frogs (N = 33 mild origin, N = 28 harsh origin) were transferred from the laboratory to environmental chambers to begin acclimation for overwintering. The temperature in each chamber was reduced at a rate of 1.5°C per day (24h), until reaching final experimental temperatures, 2°C (unfrozen treatment), and −2°C (frozen treatment). Frogs from harsh origin winter environments and exposed to frozen (−2°C) artificial winter treatment were considered to be in “local” conditions because their origin winter environments are, on average, below freezing. Likewise, frogs from mild origin winter environments and exposed to unfrozen (+2°C) artificial winter treatments were considered to be in “local” conditions because their origin winter environments are, on average, above freezing. Harsh-origin frogs in the unfrozen artificial winter treatment, and mild-origin frogs in the frozen artificial winter treatment were considered to be in “foreign” winter conditions. While in the environmental chambers, frogs were maintained individually in 16-oz. plastic cups lined with a moist paper towel, stocked with leaf litter, and covered with a screened lid. Frogs were fed at the start of acclimation, and at day five of acclimation (14.5°C), after which they ceased feeding and were no longer offered fruit flies.

### Sample collection

Nine days after reaching the final experimental temperatures, frogs were removed from artificial winter conditions and euthanized to collect samples. This time point is the longest time period juveniles have been shown to survive freezing with 100% survival (Layne Jr. et al., 1998). Frogs were euthanized via immersion in MS-222 and then double-pithing. Frogs were quickly dissected on a pre-frozen silicone dissection mat to collect ventral skin. Skin samples (ca. 25mm^2^) were stored in RNAlater preservation buffer, kept at 4°C for 24h and transferred to - 80°C for storage until RNA extraction. Entire livers were collected immediately after ventral skin; liver tissue was frozen on dry ice and then transferred to −80°C for storage until they could be dried and weighed. We also sexed animals during dissection and noted the presence and number of fat bodies visible in the abdominal cavity. Carcasses (*sans* liver and a piece of ventral skin) were frozen for dry weight and lipid extraction. Only samples from male frogs were used for analyses presented herein.

### Energy storage phenotype

We measured the amount of ingested energy that frogs stored as liver reserves by removing the whole liver, drying it at 50°C overnight and weighing it to 0.00001 g (0.01 mg). We also measured the dry mass of the frog carcass *sans* liver to the nearest 0.01 mg. To estimate the amount of ingested energy that frogs stored as lipid, we extracted lipids from dried frog carcasses following the methods of Bevier (Bevier, 1997). Dried frog carcasses were weighed to the 0.00001 g, then soaked for 24h in petroleum ether to remove triglycerides, then dried again to a constant mass at 50°C. We repeated the procedure so that each carcass underwent two 24h cycles of solvent immersion. We determined that two cycles of immersion were sufficient to extract most lipids using positive controls with known lipid fractions. For both measurements, initial (different from described) methods for estimating liver and lipid fractions of stored energy were unsuccessful. Samples that were processed using initial, unsuccessful methods were excluded from analysis; we present results for liver and lipid stores from the remaining 67 samples.

### RNA extraction, library preparation and sequencing

We extracted RNA from skin tissue using the RNeasy Mini Kit (Cat #74104, Qiagen, Hilden, DE) following the manufacturer’s protocol. We included an on-column DNase digestion step to improve the purity of the RNA yield. Extracted RNA was quantified using the Quant-it RiboGreen RNA Assay Kit (Cat # R11490 ThermoFisher, Waltham, MA, US). Prior to library preparation, RNA from each sample was diluted to equal concentration (2ng/μL). We used the Mercurius BRB-seq library preparation kit (Cat # PN 10813, Alithea Genomics, Epalinges, CH) and followed the manufacturer’s protocol to prepare mRNA libraries for Illumina sequencing. Briefly, we reverse transcribed each RNA sample with barcoded oligo-dT primers. After reverse transcription, we created a small library by pooling 5μL of each barcoded samples one tube. This library was purified, then free primers were digested, and second strand cDNA was synthesized. Full-length cDNA was then tagmented using a Tn5 transposase preloaded with adapters for library amplification. After tagmentation, the 5’ terminal fragments in the library were indexed with a Unique Dual Indexing adaptor and amplified for 15 PCR cycles. The indexed library was then purified with magnetic beads, and a small aliquot was used for quality control (Qubit, and Tapestation fragment analysis). To ensure that each sample would be represented in equal proportion in our final library, we first created a low-input library which we sequenced at a shallow read depth. All steps performed to create the low-input library for shallow sequencing were then repeated to create the final library, however we used up to 15μL of reverse transcription product to create the final library (rather than 5μL).

After shallow sequencing of our small library (NovaSeq PE 100 cycle) we demultiplexed the reads and generated read counts from each sample (see bioinformatics details below). We calculated the fold change in reads generated between each sample, and the sample with the most reads (for example, the fold change between the sample with the most reads and itself was 1, and the fold change between the sample with the most reads and a sample with half as many reads was 2). We then adjusted our input of reverse transcription product for the full library preparation to reflect these differences in concentration, to improve the evenness of each sample’s representation in the final library. Finally, we sequenced our larger library to one billion total paired end reads on a NovaSeq SP (v1.5 reagent kit) 300 cycle run.

### Bioinformatics

To generate read counts from raw sequencing data, we used Trimmomatic (Bolger et al., 2014) to trim R1 reads to 28 bp to retain the 14 bp barcode and 14 bp UMI for each sample (all genomic sequencing data were contained in the R2 read). Then, following the Alithea Genomics User Guide: BRB-seq Library Preparation kit (https://25700946.fs1.hubspotusercontent-eu1.net/hubfs/25700946/MERCURIUS-BRBseq-kit-PN-10813-10814-11013-11014-User-Guide-v0.7.1.revB.pdf) we used the “solo” mode in the STAR aligner (Dobin et al., 2013) to simultaneously align reads and produce read count matrices. Specifically, we aligned R2 reads to the wood frog genome (GenBank Accession: JAQSEC000000000.1) and used barcode and UMI data from R1 to demultiplex and deduplicate samples and read counts, respectively. Annotations were prepared according to the methods in ref et al.. Briefly, we annotated peptides from our data with the EnTAP tool (v. 0.9.0) (Hart et al., 2020) which compares peptide sequences from sequencing data against peptide sequences in three reference databases, the RefSeq complete protein database, the UniProt SwissProt database, and the NCBI NR database and uses the eggNOG mapper to assign GO terms within the EnTAP pipeline (Huerta-Cepas et al., 2016).

The eggNOG mapper for GO term annotation identifies orthologs to query sequences in the eggNOG database (Huerta-Cepas et al., 2016), which is a phylogenetically informed database of orthologs and functional annotations. eggNOG-mapper transfers GO terms from identified orthologs to query sequences, dynamically adjusting the taxonomic scope from which GO terms are transferred for each query.

### Statistics

To assess whether frogs from different origin winter environments varied in liver and lipid reserves, we used Bayesian mixed-effects models to predict either liver or lipid mass as a function of a frog’s scaled dry mass, origin winter environment, artificial winter treatment, and the interaction between origin environment and treatment. We included random intercepts for lipid extraction batch and population for the model of lipid percent, and a random intercept for population for the model of liver. Models were constructed using the ‘stan_lmer’ function from the *rstanarm* package in R (Goodrich et al., 2023) using default (weakly informative) priors and four chains for 4000 iterations after a 200 iteration burn-in period. We checked for chain convergence by examining caterpillar plots and confirming that *Rhat* statistics were all < 1.1; we report the 95^th^ highest posterior density credible intervals from the posterior sampling distribution.

We imported a matrix of unique, deduplicated read counts per sample for each gene into R (v. 4.2.2) for differential gene expression and other analyses. Prior to statistical analysis, we filtered genes for low expression (fewer than 50 reads across all samples) and filtered samples for low read counts (fewer than 10,000 reads across all genes). We analyzed differential gene expression using the DESeq2 package in R (Love et al., 2014). Differential expression was determined using the following model:

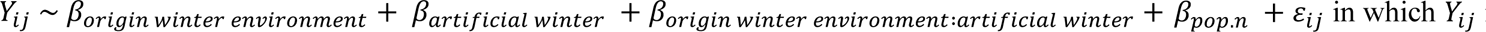 is a count of transcript *i* in individual *j*. *β*_*origin winter environment*_ is a fixed effect with two levels, mild and harsh, denoting whether a frog came from an origin winter environment that was short and warm or long and cold, respectively. *β*_*artificial winter*_ is a fixed effect with three levels corresponding to whether a sample was collected from a frog before artificial winter treatments commenced (pre), or from a frog in unfrozen (+2°C), or frozen (−2°C) artificial winter treatment. *β*_*pop.n*_ is a nested random effect within the fixed effect “Origin winter environment” with three levels (1,2,3) representing the three populations per origin winter environment sampled.

To visualize how patterns of gene expression varied among populations and treatments, we clustered the gene expression data using principal components analysis of the variance-stabilized transformed counts of the 500 most variable genes. Following the methods used by Kenkel and Matz (2017), we used discriminant analysis of principal components (DAPC) using the *adegenet* (Jombart et al., 2010) package in R to plot the maximal distances on the first linear discriminant between groups of frogs in their local winter conditions. Onto this axis, we projected gene expression profiles of frogs in foreign winter conditions. We compared the absolute distances between same-origin frogs in local and foreign winter conditions using (default) weak priors and the MCMCglmm function from the *MCMCglmm* (v. 2.35) package in R (Hadfield, 2010).

We identified groups of genes (modules) whose changes in expression varied similarly in response to variables in our dataset using weighted gene co-expression network analysis (WGCNA) using the *WGCNA* (v. 1.72-1) package in R (Langfelder & Horvath, 2007). For WGCNA analysis, we also searched for gene modules that were expressed in association with the mass of liver reserves, lipid reserves, and binary terms for whether frogs were exposed to foreign or local winter conditions. WGCNA modules were constructed using a soft thresholding power of three and a minimum module size of 100 genes/module). After module construction, we used the ‘lmFit’ and ‘eBayes’ functions from the R package *limma* (v.3.54.2) to model the expression of each module’s eigengene in response to the same explanatory variables used in DESeq2 analysis (additive effects of origin winter environment, artificial winter treatment, and their interaction) (Ritchie et al., 2015). Detailed methods for WGNCNA are presented in the supplemental methods.

We used the ‘goseq’ (v 1.5.0) package (Young et al., 2010) in R to identify enriched gene ontology (GO) terms in lists of differentially expressed genes from DESeq2 analysis, and in lists of genes belonging to modules identified by WGCNA as having the most significant correlations with experimental variables. Importantly, differentially downregulated gene sets can be enriched for GO terms; in these cases, GO terms are significantly overrepresented among genes with significant negative log2fold changes. In several visualizations we highlight terms descended from the immune system process (GO:0002376), innate immune response (GO:0045087), and the adaptive immune response (GO:0002250) parent GO terms.

## Results

Liver but not lipid reserves varied significantly between frogs from different origin winter environments. We saw a significant effect of origin winter environment on the amount of liver reserves, in which frogs from places with mild winters had smaller livers in all conditions (controlling for body size) than frogs from places with harsh winters (β_MildOrigin_ = −1.2 mg, 95% HPD interval [-2.1 mg,-0.47 mg]). Similarly, liver reserves varied significantly in response to an interaction between origin winter environments and artificial winter treatment (Figure 2). Compared to the timepoint before exposure to artificial winter treatment (pre), liver reserves were greater in the unfrozen (+2°C) treatment for harsh origin frogs, and lower in the unfrozen (+2°C) treatment for mild origin frogs (β_MildOrigin:UnfrozenTreatment_ = −1.1 mg, 95% HPD interval [−2.1 mg, −0.04 mg]). Regardless of origin winter environment, liver reserves were greater in frogs that were larger (for each standard deviation increase in frog dry mass, liver reserves were 1 mg, 95% HPD interval [0.78 mg, 1.3 mg] larger). Lipids comprised a very small percentage of the dry mass of wood frogs in our study (β_Intercept_ = 2.5 mg, 95% HPD interval [1.6 mg, 4.1 mg]). Like liver, lipid reserves were greater in larger frogs (each standard deviation increase in frog size corresponded to a 1.2 mg (95% HPD interval [0.8 mg, 1.5 mg]) increase in lipid reserves). Lipid reserves did not vary significantly with origin winter environment, artificial winter treatment, or their interaction (all 95% HPD intervals contained 0) (Figure 2). Data on wet and dry masses of frog carcasses, livers, and lipids are provided in table S1.

**Figure 2.**
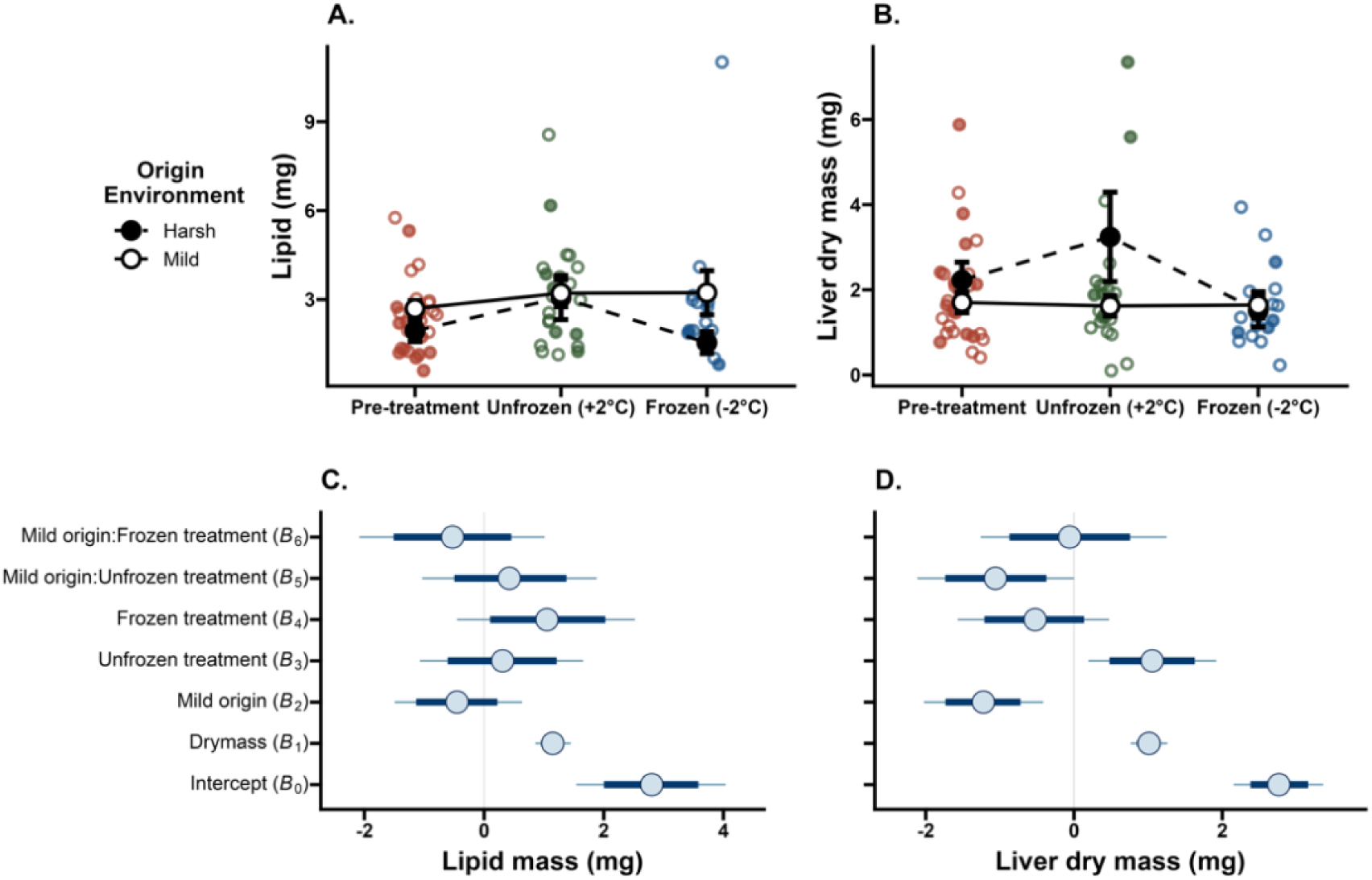
Harsh origin frogs (filled circles) and mild origin frogs (open circles) maintained similar proportions of lipids in all conditions (pre-winter, unfrozen winter (+2°C), and frozen (−2°C) winter). Liver dry mass was higher among frogs from harsh origin winter environments in pre-winter, and unfrozen winter conditions, but did not differ from mild origin frogs in the frozen winter conditions.

We also detected patterns at the transcriptomic level related to energy stores and metabolism. Despite finding no differences in gross lipid proportion between populations or among treatments in harsh-origin frogs, genes that were downregulated in freezing compared to pre-winter treatments were enriched for many lipid metabolic processes, including lipid catabolic process, fatty acid oxidation, and lipid oxidation (Figure 3; FDR_all_ < 0.05; table S2 for lists of significantly enriched GO terms). In contrast, no biological processes were enriched among genes that were significantly downregulated in freezing versus pre-winter treatments in mild-origin frogs. In freezing (−2°C) artificial winter conditions, frogs from mild origin winters were enriched for the lipid droplets GO term (a cellular component, GO:0005811) compared to frogs from harsh origin winters (FDR < 0.01). In the same contrast (mild origin versus harsh origin, in −2°C conditions), mild origin frogs were also enriched for the short-chain fatty acid catabolic process (a child term of lipid metabolic process; GO:0019626, FDR = 0.08).

**Figure 3.**
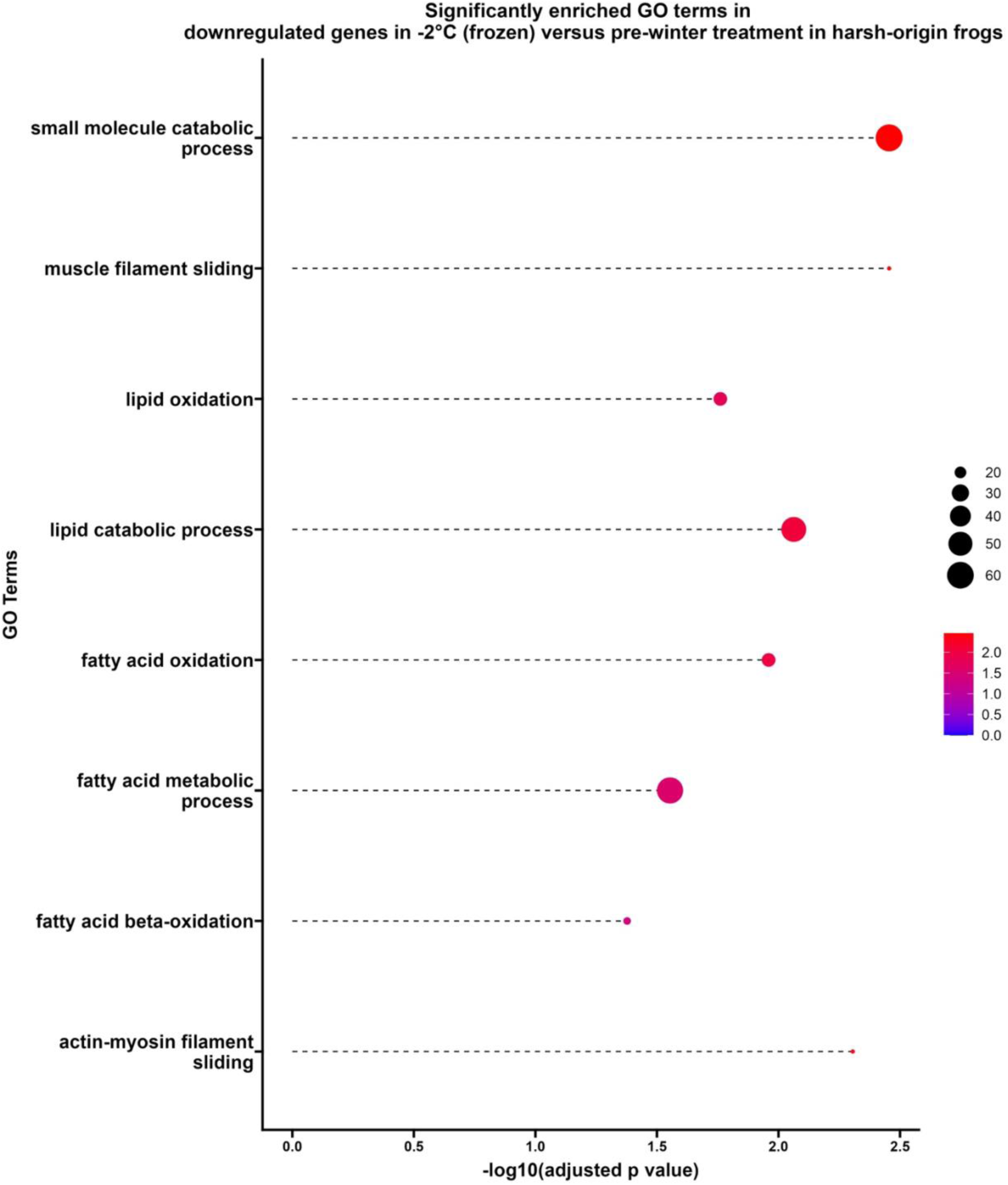
As predicted, biological processes associated with fatty acid oxidation, fatty acid metabolic processes, and lipid oxidation were overrepresented among significantly downregulated genes in frogs in the freezing (−2°C) treatment. Figure shows results for harsh-origin frogs but results are similar for mild-origin frogs. Larger bubbles correspond to GO terms with better representation in the dataset, and warmer (more red) colored bubbles indicate more significant adjusted p values.

Much of the differential gene expression we measured was due to either a main effect of origin winter environment or artificial winter treatment (Figure 4, Figure 5). In principal components space, the first axis (17% variation explained) separated points based on artificial winter treatment, while the second axis (12% variation explained) separated points according to origin winter environment. The identities of differentially expressed genes varied markedly with origin winter environment, but gene expression overall showed similar amounts of plasticity between origins. We estimated plasticity by measuring the absolute distance between frogs in local versus foreign environments on the first axis of a linear discriminant function derived by maximizing variation between frogs in their native environments (arrows on Figure 6). In “foreign” winter conditions, frogs from harsh origin environments became more like frogs from mild origin environments, and vice versa. Moreover, reciprocally exposing frogs to foreign artificial winter treatments (−2°C for mild-origin frogs and +2°C for harsh-origin frogs) was associated with similar magnitudes of change (plasticity) in their overall gene expression profiles (*P* = 0.99; Figure 6). Exposure to cold (+2°C or −2°C) influenced gene expression, and placement in a foreign artificial winter treatment drove gene expression profiles toward the local expression profiles of the contra-origin frogs. However, regardless of cold-exposure or local/foreign exposure status, frogs still grouped primarily based on origin winter environment (Figure 6).

**Figure 4.**
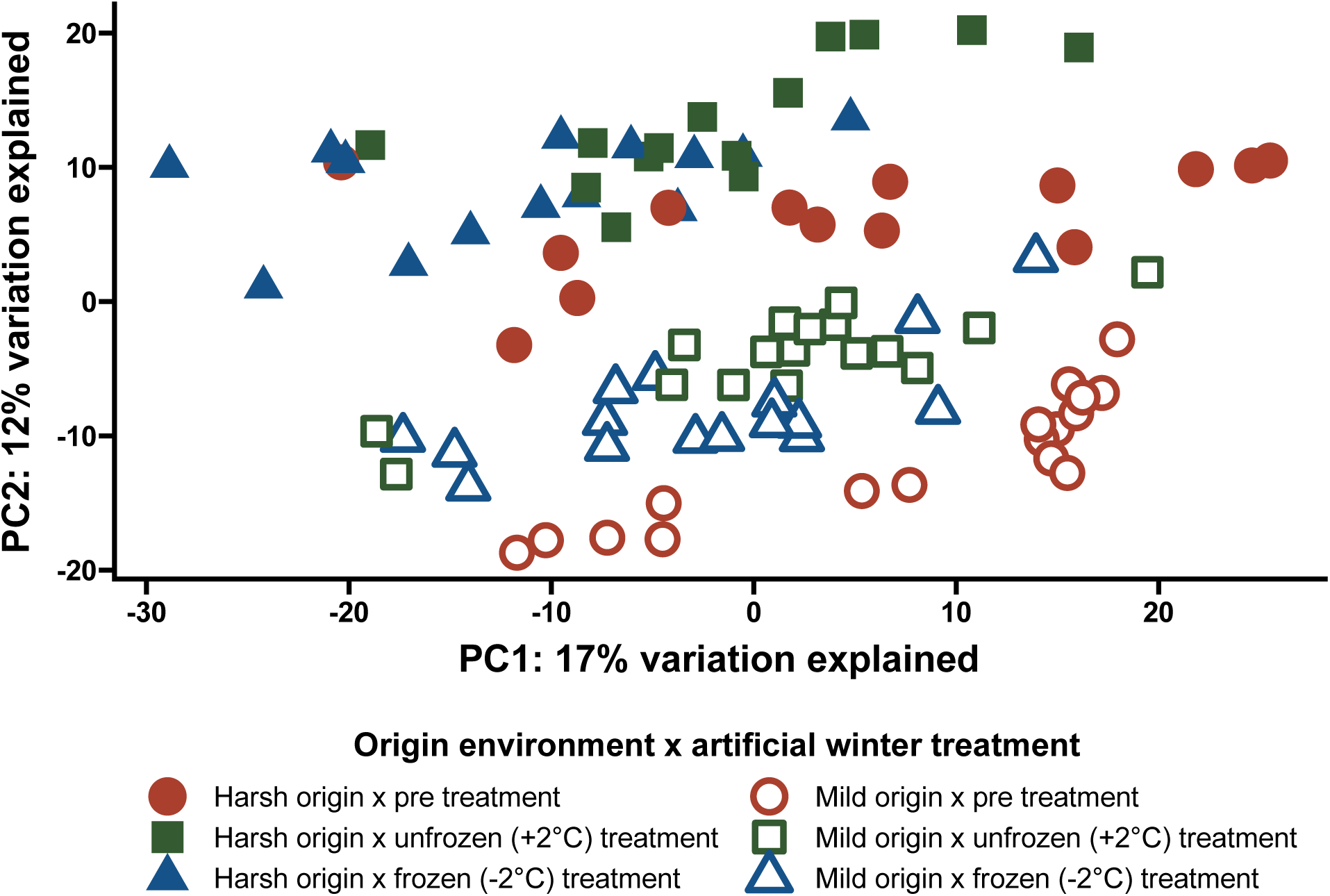
Ventral skin gene expression varied by both artificial winter treatment (PC1) and frog origin (PC2). We examined, but did not plot PC3 (9% variation explained), which also separated samples by frog origin. Pricincipal coordinates were calculated the variance-stabilized transformed values of the 500 most highly variable genes.

**Figure 5.**
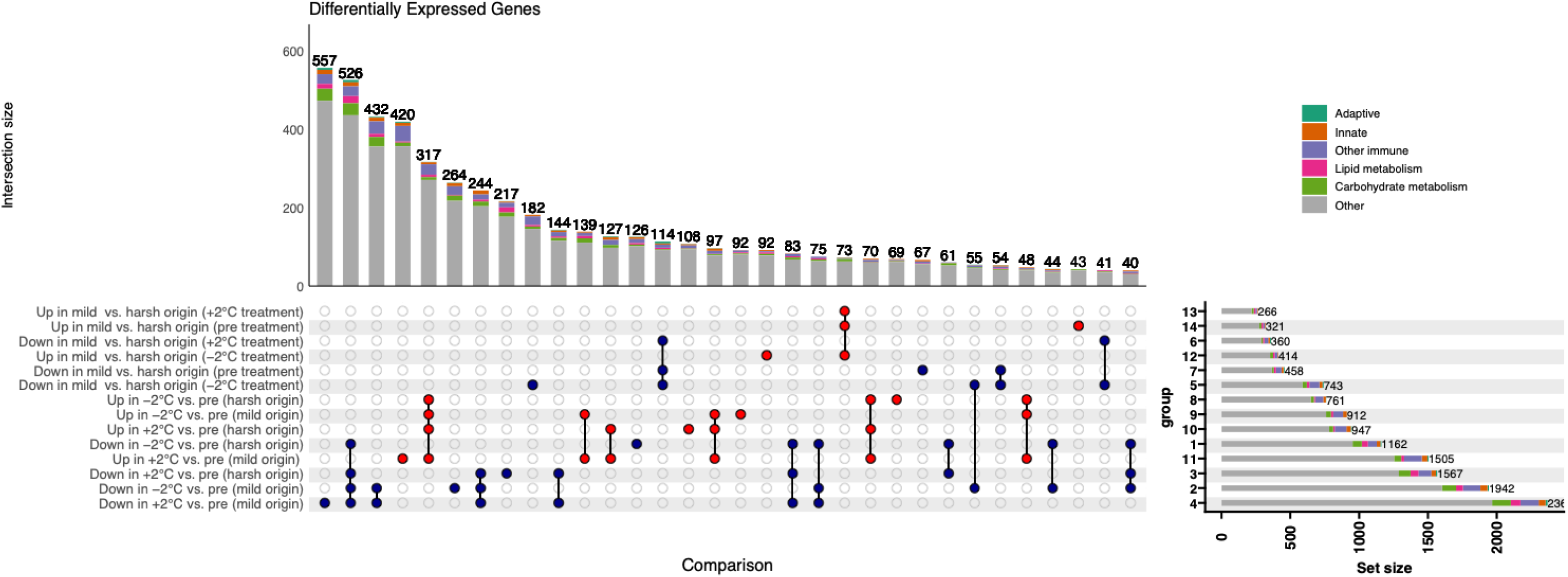
UpSetR style figure depicting differentially expressed (FDR < 0.1) genes. Written descriptions of each statistical comparison are shown at the left, separated by the direction of log2fold expression change (red – positive/up, blue – negative/down). The size of gene sets is shown in the right-hand bar graph, indicating the total number of differentially expressed genes for each comparison and direction. Intersections (genes that are differentially expressed in multiple comparisons) are shown with colored circles connected with black lines. Intersection set size—the number of gene differentially expressed in multiple comparisons— is shown in the top bar graph. Genes annotated with parent terms of the following processes are indicated in color: adaptive immune response parent term (GO:0002250) in dark green; innate immune response parent term (GO:0045087) in orange; immune system process parent term (GO:0002376) (but not with either the adaptive or innate parent terms additionally) in purple; lipid metabolic process (GO:0006631) in pink; carbohydrate metabolic process (GO:0005975) in bright green. Genes annotated with all other GO terms are shown in gray. Note that a minimum threshold of 40 genes per intersection set was enforced for this visualization, so some comparisons and intersections are not visualized.

**Figure 6.**
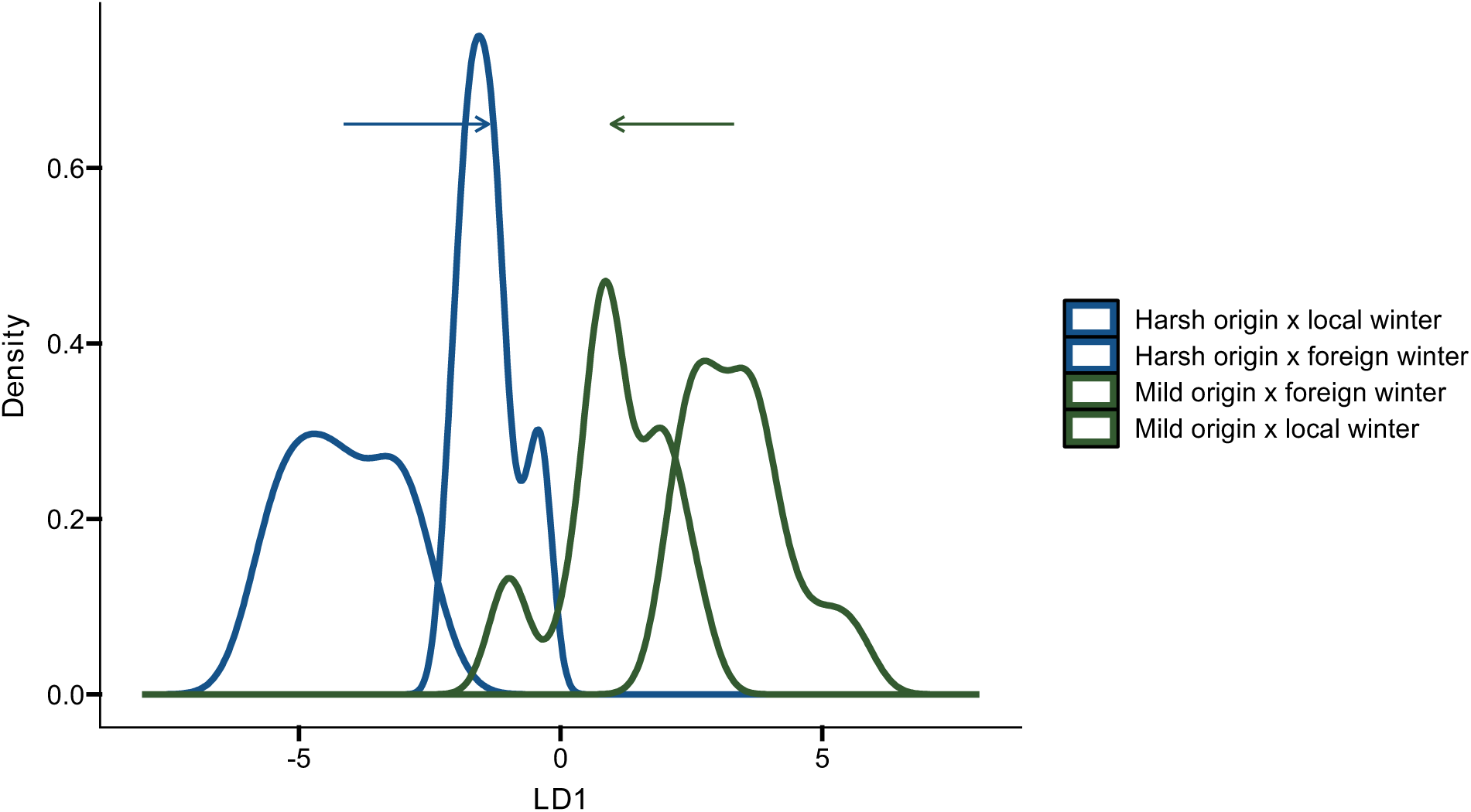
Frogs from harsh and mild winter environments express similar levels of plasticity in their gene expression profiles between local and foreign winter environments (*P*_MCMC_ = 0.98). The x axis (LD1) is a discriminant function that maximizes the difference between harsh-winter (blue-outline) and mild-winter (green-outline) frogs in their native environments (color-filled curves). Gray-filled curves represent frogs in foreign winter conditions (harsh winter frogs in unfrozen (+2°C) conditions, and mild winter frogs in frozen (−2°C) conditions). Arrows above curves indicate the mean MCMC posterior estimate for the difference in gene expression profiles between frogs in their native winter conditions and frogs in their reciprocal (foreign) winter conditions.

We found a small number of genes (15) that were differentially expressed (FDR < 0.1) in response to an interaction between the origin of a frog and the artificial winter treatment to which a frog was exposed. Of these, two were unannotated, and two were not annotated with a predicted gene identity (but were annotated with a description). Of the 13 genes that were annotated with a predicted gene identity, two were differentially expressed in multiple interactions. In mild-origin frogs, expression of the *HSPA8* (heat shock protein family A (Hsp70) member 8) gene was dramatically increased in above freezing (+2°C; FDR = 0.04) and freezing (−2°C; FDR *>* 0.01) artificial winter treatments compared to the pre-winter treatment. In the pre-winter treatment, however, *HSPA8* was expressed at similar levels in frogs from both origin environments. Conversely, in harsh-origin frogs, expression of *HSPA8* remained very low across treatments (Figure 7A). Another interaction was apparent in the expression of the gene intelectin 1, *ITLN1*. In mild origin winter frogs *ITLN1* was expressed at stable, relatively low levels in all tested treatments. In harsh-origin frogs, however, expression of *ITLN1* was expressed at low levels, indistinguishable from mild-origin expression levels, in the frozen artificial winter treatment, and significantly downregulated compared to the pre-winter (FDR<0.001) and unfrozen artificial winter treatments (FDR = 0.1; Figure 7B). Although on average, *ITLN1* expression was higher in harsh-origin frogs, expression in this group of individuals was marked by a large amount of variability. The spread in expression levels for this gene in the harsh-origin unfrozen treatment frogs was not explained by differences amongst the three populations comprising the harsh-origin frogs. In mild-origin frogs, expression of the secretory leukocyte peptidase inhibitor gene (*SLPI)* monotonically decreased from pre-to +2°C, to −2°C treatments, while in harsh-origin frogs, expression was significantly upregulated in both unfrozen and frozen treatments, compared to pre-winter frogs (FDR_both_ < 0.0001; Figure 7C). Finally, the IFNγ receptor 2 (*IFNGR2*) gene was significantly downregulated at the pre-winter timepoint in mild-origin frogs compared to harsh-origin frogs. This pattern was reversed such that mild-origin frogs expressed *IFNGR2* at higher rates than harsh-origin frogs in the freezing artificial winter treatment (FDR = 0.02; Figure 7D). All genes that were annotated with a predicted gene identity or a description and differentially expressed in an interaction between origin and artificial winter treatment are presented in Table S3.

**Figure 7.**
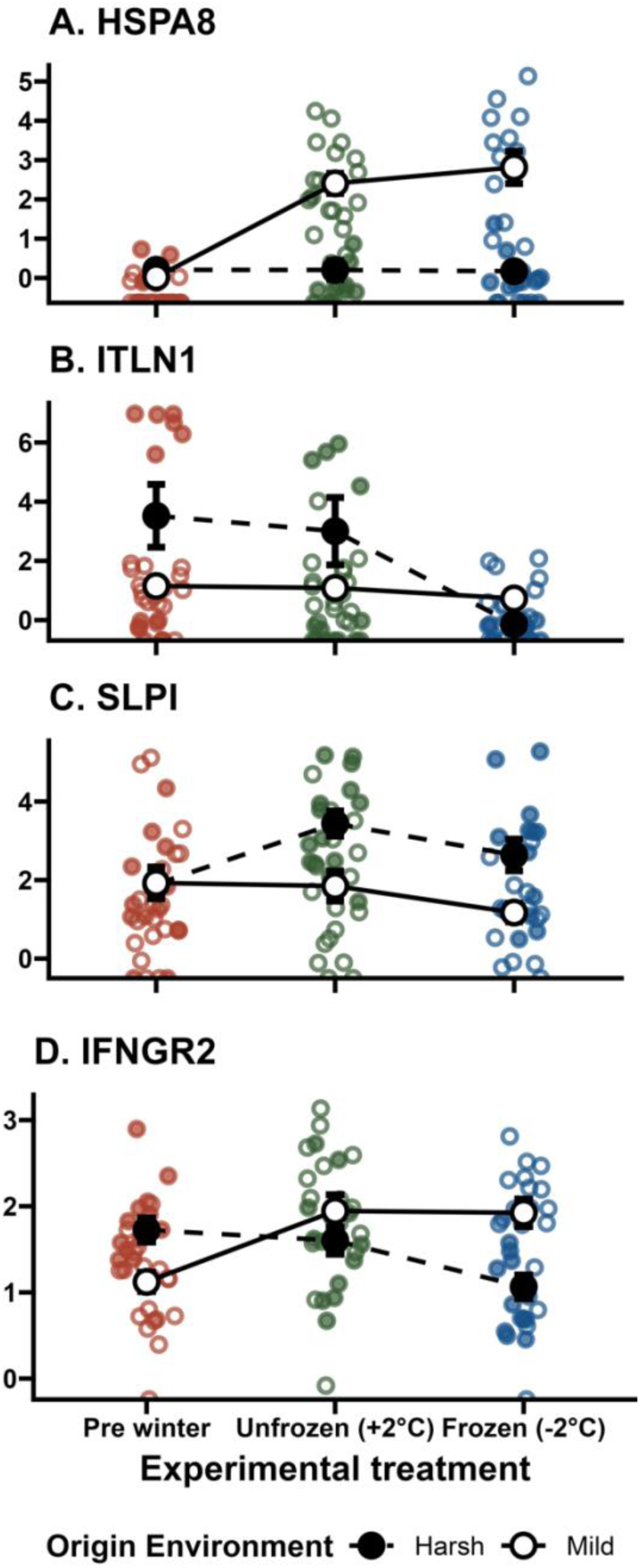
Expression (log10 values of normalized counts) of four genes—*HSPA8, ITLN1, SLPI,* and *IFNGR2—*differentially expressed in response to the interaction between frog origin environment and artificial winter treatment.

Across origins, cold exposure (both +2°C and −2°C) resulted in coordinated gene upregulation (differential gene expression that was dominated by enrichment of GO term processes) but not coordinated downregulation. By count and across both origin winter types, exposure to cold conditions (+2°C or −2°C) resulted in a greater number of shared significantly differentially downregulated (FDR < 0.1 and log2fold change <0) compared to upregulated genes (log2fold change > 0; 513 down vs. 284 up; Figure 5). Despite being fewer, genes significantly upregulated in both cold treatments across origins were enriched for 81 GO terms indicating that cold-induced upregulation is likely a conserved, coordinated process. The biological processes enriched in the cold included the GO term for the cellular response to stress (FDR < 0.05; GO:0033554) and the term for apoptotic processes (FDR < 0.05; GO:0006915). This coordinated enrichment of many biological processes in genes that were upregulated in cold conditions regardless of frog origin, was not seen in the larger number of downregulated genes. In cold artificial winter conditions, neither mild-nor harsh-origin frogs had any shared enriched GO terms among significantly downregulated genes.

One of the most striking differences between frogs from different origins was the constitutive downregulation of immune processes in frogs with a mild origin. In all treatments (pre-winter, - 2°C and +2°C), frogs from mild origin environments significantly downregulated genes annotated with the GO terms for immune effector processes (GO:0002252) and regulation of adaptive immune response (GO:0002819). Comparing frogs from different origin environments on a pairwise (treatment-by-treatment) basis revealed that many additional immune-related processes were downregulated in frogs with a mild origin. For example, in the freezing (−2°C) artificial winter treatment, frogs with a mild origin winter significantly downregulated many biological processes related to adaptive immunity and antigen processing. In unfrozen artificial winter treatment (+2°C), processes associated with cell activation, leukocyte and lymphocyte activation, defense response to viruses, and adaptive immune responses were enriched among significantly downregulated genes in mild origin frogs compared to harsh origin frogs (Table S2). In addition, 37 of the 51 enriched GO terms in the Green module from WGCNA analysis, which included genes upregulated in frogs from harsh origin winter environments, and downregulated in frogs from mild origin winter environments, were associated with immune processes (Figure 8; Figure 9). We examined whether frogs from different origin winter environments were similarly plastic in their expression of genes in the Green module. In contrast to global expression patterns (considering all genes), we found that harsh-origin frogs expressed genes in the Green module in a more plastic manner (P = 0.0003) than mild-origin frogs (Figure 10).

**Figure 8.**
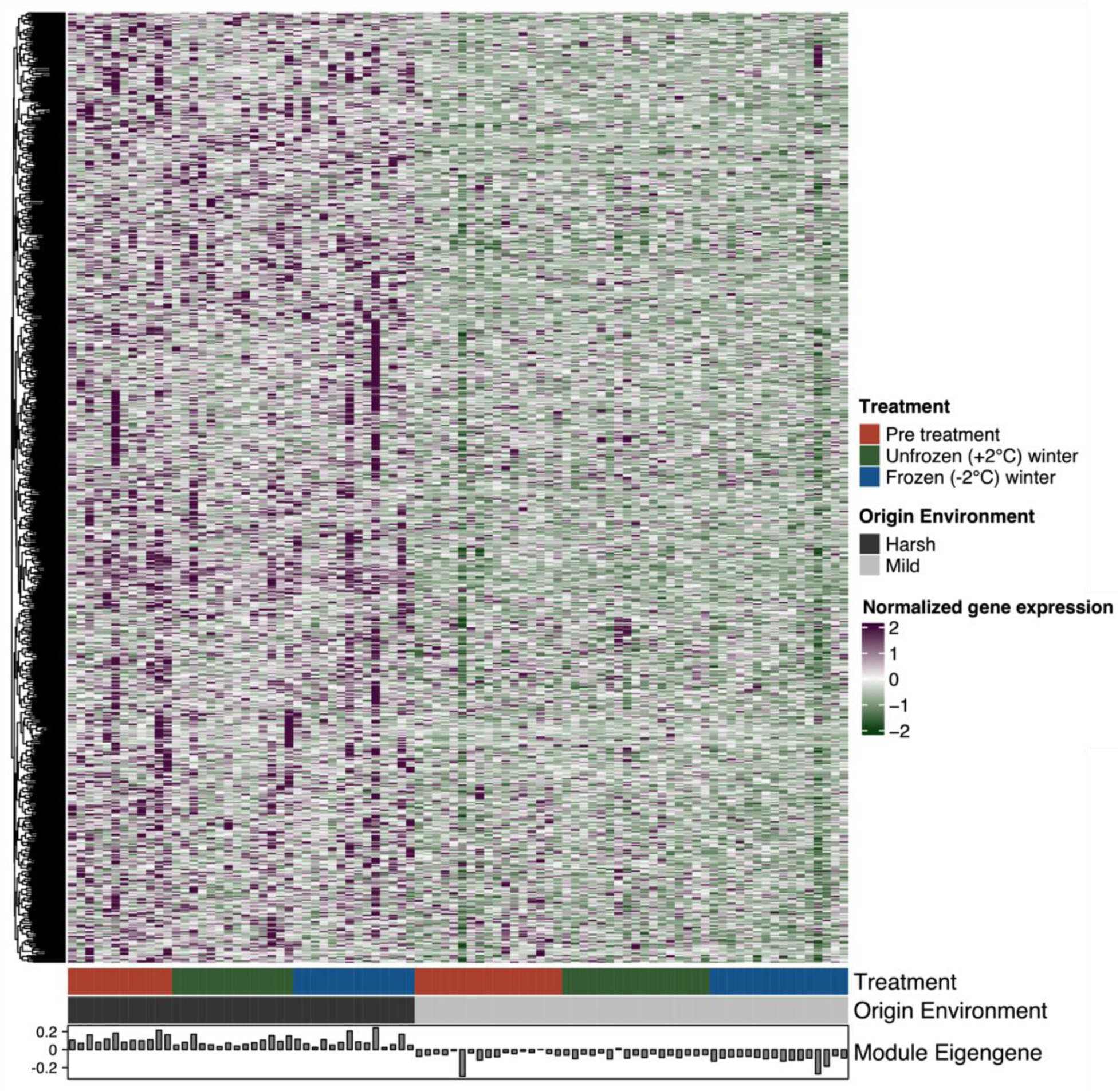
A major difference between frogs from different origin environments was constitutive upregulation of immune processes. Of the 13 gene modules identified from weighted gene co-expression network analysis, the green module (which contained 751 genes) was highly associated with origin environment (*Pearson’s R =* 0.86, FDR_Green eigengene_ < 0.0001). Genes in the green module were overwhelmingly upregulated (purple/darker shade) in frogs from harsh origin environments and downregulated (green/lighter shade) in frogs from mild origin environments. GO term enrichment analysis on this module identified 53 enriched GO terms among upregulated genes, and of those, 37 (70%) were related to immune functions.

**Figure 9.**
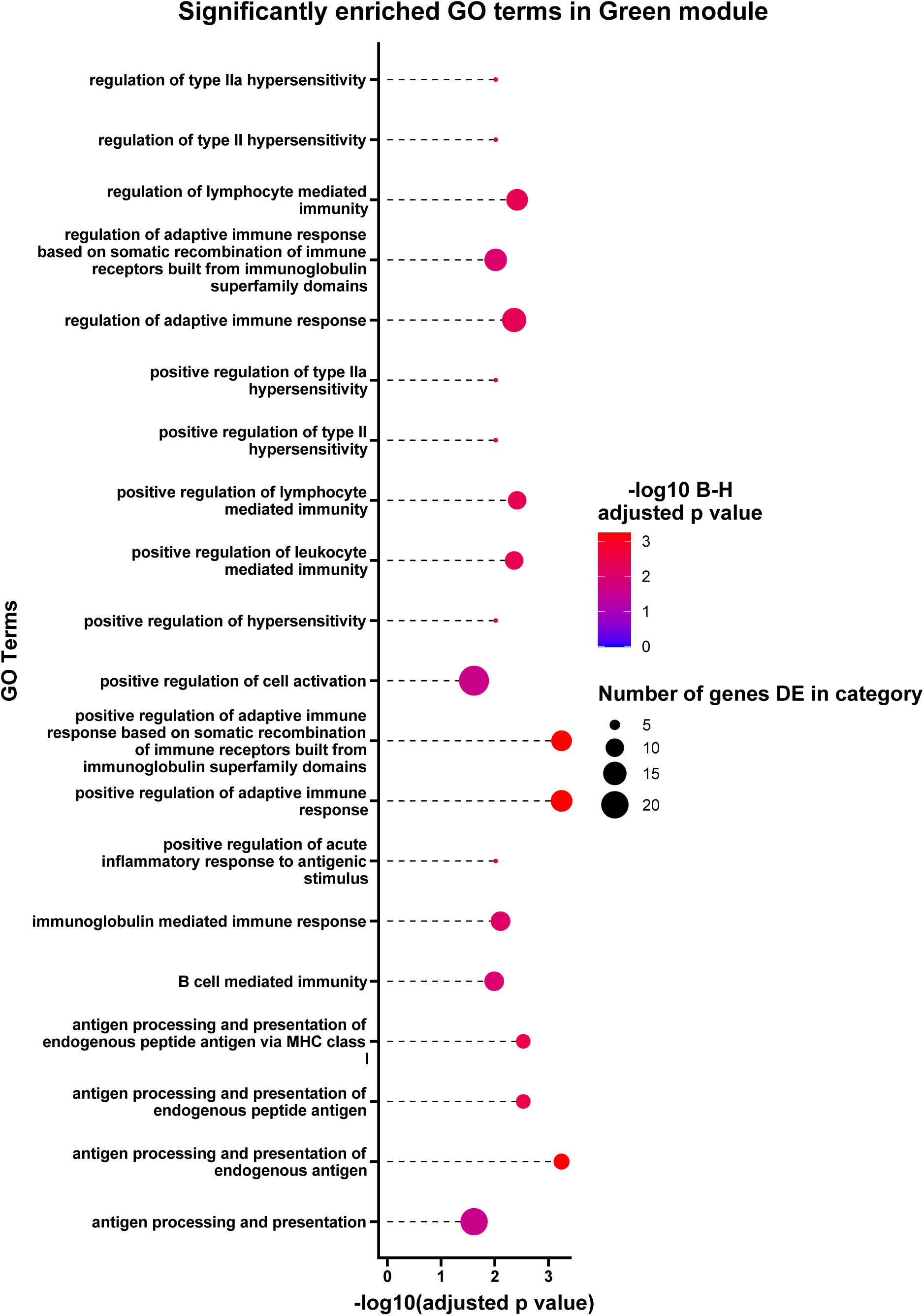
Biological processes related to immunity dominated (comprised 70%) of the enriched GO terms in the Green module, in which gene expression was upregulated among harsh-origin frogs and down-regulated among mild-origin frogs.

**Figure 10.**
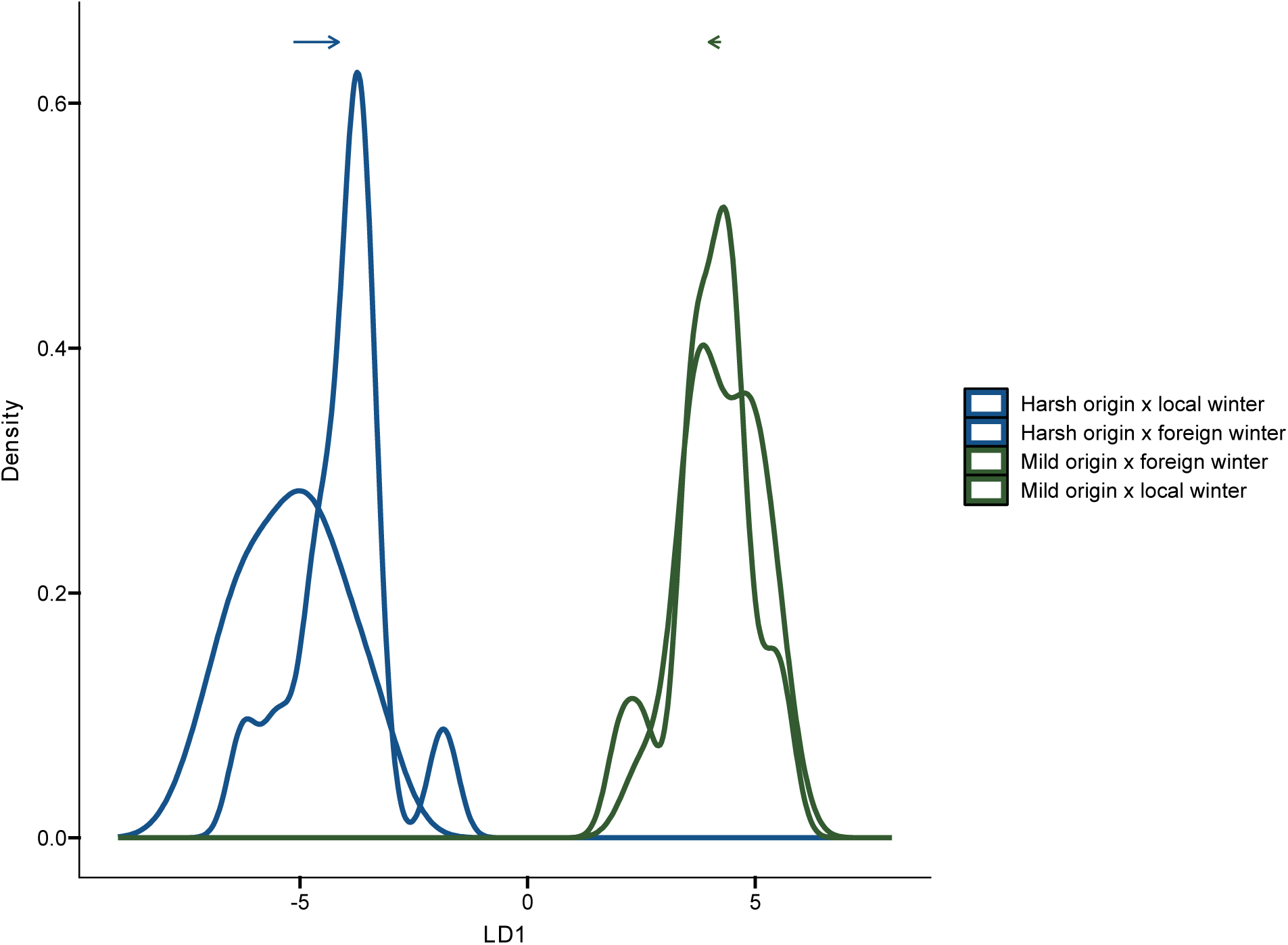
Expression of the 751 genes from the Green module, which were constitutively upregulated in harsh-origin frogs was significantly (p = 0.0003) more plastic in harsh-origin frogs than in mild-origin frogs.

## Discussion

This research highlights how winter climates have shaped the evolution of wood frog populations. We call attention to one avenue by which winter warming may drive intraspecific variation: via effects immune gene expression. In addition, we tested the hypothesis that different energy storage phenotypes among frog populations comprise the mechanistic basis for the relationship between ancestral winter climates could drive immune gene expression patterns. We found that ventral skin expresses distinct transcriptional profiles in unfrozen (+2°C) and frozen (−2°C) overwintering temperatures, and that gene expression in this tissue varies markedly with the genetic origin of the population. Frogs from places where winters have historically been colder and longer did store proportionally more ingested energy in the liver when unfrozen and upregulate several immune processes in all tested conditions as compared to frogs from mild origin winter environments.

Examining whether local adaptations to winter conditions contribute to variation in immunity across populations is particularly relevant given changing winter climates, susceptibility of wood frogs to infectious diseases, and the status of wood frogs as an emerging model for comparative amphibian immunity (Crespi et al., 2015; Douglas & Katzenback, 2023; Eskew et al., 2018). A recent winter climate modeling effort found that future winter energy requirements will likely be reduced across much of the wood frog range (M. J. Fitzpatrick et al., 2020). Acting alone, energetically less-expensive future winters could prove beneficial to wood frogs, but the multifaceted nature of winter climate change, and the expansive geographic distribution of wood frogs mean that the magnitude and nature of wood frog responses to changing winters are likely to vary on a population-by-population basis. In particular, the importance of winter as a selection pressure on energy allocation strategies may wane as winters become shorter and warmer, and could be surpassed by other challenges, such as infection risk.

We reared frogs for the same amount of time before exposing them to artificial winter temperatures. Assuming equivalent developmental rates, if frogs from different origin winter environments start the winter with different ratios of stored carbohydrates and lipids, this result could be evidence that growth and energy accumulation strategies are adapted to local winter conditions. Our findings, that liver size varies with origin winter environment, agree with previous research showing differences in liver and glycogen reserves between Alaskan and Ohioan wood frog populations (Costanzo et al., 2013). Frogs from populations that have historically tolerated freezing more frequently and for longer tend to have larger livers when adjusted for body size. The significant interaction we saw, in which liver size was greater in harsh-origin frogs (as compared to mild-origin) in pre-winter and unfrozen (+2°C) artificial winter treatment, but not higher in the freezing (−2°C) artificial winter treatment also makes sense. While we expected frogs from harsh-origin environments to commit more accumulated energy to liver reserves prior to winter and to maintain these reserves prior to the onset of freezing, we did not expect to see this pattern in the mild-origin frogs. When exposed to freezing conditions, however, we expected liver reserves to decrease in all frogs (Costanzo et al., 2015), and indeed this decrease is what we observed.

Unlike previous comparisons of Alaskan and Ohioan frogs (Costanzo et al., 2013), we found no differences in lipid accumulation between frogs from different origin winter environments. Notably we restricted regional comparisons to population that exist within a much smaller geographic area, all occurring within the Eastern mitochondrial DNA clade of wood frogs (Lee-Yaw et al., 2008). Differences in lipid accumulation, or the lack thereof, may be attributable to the magnitude of local adaptation to winter climates, or to the differing effects of drift, and physical and chronological separation among populations and clades. Additionally, most previous studies documenting differences in accumulated energy reserves have often used wild-caught adults subjected to acclimatization periods in the laboratory rather than frogs reared from egg masses (e.g., Costanzo et al., 2013). In studies of energy accumulation, this approach may suffer from lingering plastic responses to prey availability prior to capture, which may also be confounded with origin winter environment. By rearing animals from the egg stage in identical laboratory conditions, our study circumvents this complication. An alternative possibility is that the method we used to measure differences in lipid accumulation, solvent extraction, was not sufficiently sensitive to detect meaningful differences in how frogs from different origin winter environments store and metabolize lipids. We posit this specifically because of the many differences we observed at the gene expression level between harsh- and mild-origin frogs with respect to lipid accumulation, and metabolism, all of which agreed directionally with our original predictions. A related possibility may reconcile both our finding that gross lipid reserves did not vary between frogs from different origin environments, and our finding that biological processes associated with lipid metabolism occurred at higher rates in mild origin winter frogs than in harsh. Specifically, it is possible that frogs in our study did have differences in their lipid storage and accumulation strategies at a cellular level, but that frogs were not fed an amount or variety of prey items sufficient for them to accumulate fat reserves comparable to what they would accumulate in natural conditions. Finally, unlike the clear differences in constitutive gene expression patterns between frogs from different genetic origin populations, we found less robust support for the proposed mechanism that differences in stored lipids and carbohydrates lead to different immune gene expression profiles. Because concurrent links between energy storage phenotypes and immune gene expression patterns were absent and/or weak, we suggest that other more important evolutionary and ecological correlates drive the functional relationship between ancestral winter climates and the skin immune transcriptome.

Despite our mixed findings on the storage and use of lipid reserves, our results showing regional differences in adaptations for storing and using ingested energy in the liver raise the new question of how energetic phenotypes of future wood frog populations might diverge as winters become less energetically costly? If winters no longer require freeze tolerance, will individuals with genotypes that prioritize growth over the accumulation of cryoprotectant liver glycogen, which may previously have been maladapted for harsh winters, enjoy fitness advantages (Voituron et al., 2002)?

In addition to the effects future winters will have on variation among wood frog populations and genotypes, an arguably more important question has to do with how future winters will influence the communities and competitors that wood frogs encounter. Wood frogs do best in places where other frog species can’t live; their larval stage is notably shorter than many of their congeners, allowing them to occupy vernal pools with correspondingly brief hydroperiods (Anholt et al., 2000), and their ability to tolerate freezing gives them exclusive access to higher latitudes than any other frog (Dodd, 2013). The potential advantages or disadvantages of energetically less-expensive winters at the individual organism level may be outweighed by impending changes in community structure, which are likely as winters become shorter and warmer.

Our experimental approach—a common garden and reciprocal exposure—allowed us to observe that overall plasticity in transcriptional responses of wood frogs from different origin winter environments to two artificial winter treatments was similar. Yet, we suggest that warming winters may affect frogs differently, because regardless of artificial winter conditions, gene expression varied significantly with genotype (origin winter environment). We show that wood frog populations from disparate origin winter environments express disparate immune phenotypes, before and during exposure to both frozen and unfrozen artificial winter temperatures. Further, we show that among the subset genes that are constitutively differentially expressed between frogs from different origin winter environments, plasticity in different thermal environments differed significantly. Our results indicate that future winters, which throughout most of the wood frog range are predicted to be shorter and warmer (M. J. Fitzpatrick et al., 2020), are likely to induce different effects on frogs from different origin winter environments (and different genetic backgrounds). We emphasize that we measured gene expression in a single tissue (skin) over a single simulated winter and cannot make holistic predictions of what future winters might foretell for frogs of either genetic background. We can, however, describe the nature of differences in how frogs responded to being exposed to foreign (for their genetic background) winter climates, with a focus on frogs from harsh-origin environments, which are more likely to experience milder future winters.

In general, we observed that frogs from harsh origin winter environments upregulated immune processes, including cellular immunity, inflammatory processes, and adaptive immune processes, as compared to frogs from mild winter environments. One of the genes upregulated in harsh-origin frogs as compared to mild-origin frogs when in both cold conditions was *SLPI.* This gene was also upregulated in cold temperatures in a prior study investigating changes in ventral skin gene expression throughout the course of winter in a single population of frogs from Connecticut, US (ref et al.). Notably, our previous observation of increased *SLPI* expression in cold temperatures was from a study that collected samples from frogs overwintering in semi-natural conditions (outdoors, in local leaf litter, with access to natural prey items). Combined with the expression patterns we observed, the antimicrobial properties of *SLPI*, as well as its purported immunomodulatory abilities (Majchrzak-Gorecka et al., 2016) make this gene an interesting candidate for future investigation. If increased expression of *SLPI* in cold conditions among harsh-origin frogs is adaptive, several selective pressures, such as pathology due to autoimmunity, or increased pressure from microbial pathogens could merit further study. We also observed that in unfrozen (a proxy for future) winter conditions, harsh-origin frogs down-regulated inflammatory immune processes in comparison with their expression in freezing winter conditions. Together, these results suggest that for frogs adapted to harsh winters, milder future winters might reduce immune activation, and for frogs living in places where winters have historically been mild, mild local winters may have selected for reduced immune gene expression.

We predicted that frogs from harsh-origin environments would upregulate genes associated with inflammatory immune processes while in −2°C artificial winter treatments, due to having more stored energy available, as carbohydrates would bias harsh-origin frogs toward more inflammatory cellular environments. We did find that frogs from harsh origin winter environments stored more energy in the liver (and presumably had more carbohydrates available for anerobic metabolic processes during freezing), and, in all treatments, mild-origin frogs had reduced expression of immune processes. These results provide evidence that frogs from harsh origin winter environments constitutively express higher levels of immune processes before and throughout the winter compared mild-origin frogs and leave open the possibility of a link between liver reserves and immune gene transcription. Other reasons that we saw constitutive differences in immune gene expression could have to do with parasite pressure; frogs from harsh origin winter environments may experience greater pressure from parasites in their origin environments, and higher rates of immune gene expression could be beneficial to frogs in these environments. Alternatively, frogs from mild origin winter environments may face equal or greater levels of parasite pressure as compared to those from harsh origin winter environments, but do not experience fitness benefits associated with higher levels of immune gene expression (perhaps due to immunopathology). In a 2011-2012 survey of wood frogs throughout the eastern portion of their distribution, researchers detected an extremely low prevalence of chytrid fungus infection, and a moderately high rate of ranavirus infection in the two regions where we collected egg masses for the current study (Crespi et al., 2015). Host and parasite population sizes can cycle in a frequency-dependent manner (Tompkins et al., 2001). Understanding whether wood frog populations are positioned along clines of parasite pressure, as well as tracking temporal fluctuations in parasite pressure with respect to host generations and their immune profiles would go a long way toward addressing the extent to which constitutive differences in immune gene expression profiles are directly or indirectly related to winter climate or to other ecological drivers.

Other researchers have found ecological correlates of selection on larval period and size at metamorphosis among wood frog populations across a similar geographic range (Le Sage et al., 2021). Such insights may also prove to be applicable to our understanding of the drivers of clines in immune gene expression profiles in the future. At present, however, we cannot say whether climate or microclimate characteristics, parasite prevalence, or something else is behind the large differences we observed in immune gene expression between frogs from different origin winter environments, or whether these differences are even adaptive. The preponderance of immunological processes among enriched GO terms between frogs from different origin environments, and our finding that genes in a module that was constitutively differentially expressed between origin types was significantly differently plastic, however, seem unlikely to be purely the result of drift. Further work employing population genetic methods, as well as laboratory immune challenges could resolve the potential adaptive function of these constitutive differences (Bhaskara et al., 2023).

While our study was designed to measure gene transcription in the winter, wood frogs also regulate physiological processes at post-transcriptional, translational, and post-translational levels (Al-attar et al., 2020). Thus, in cases in which we did not observe a difference in the regulation of processes related to immunity at the transcriptional level, downstream processes may still be at play, driving differences among populations, origin winter environments, and current winter conditions. Furthermore, gene expression in response to cold and below-freezing temperatures is highly tissue-specific (Chen et al., 2023; Gupta et al., 2020; Lv et al., 2024). Our findings, while limited to the ventral skin, invite further investigation such as expanding the scope of inquiry to other immune tissues such as liver, spleen, or blood, incorporating population genomic inferences, and employing infection and/or immune challenges to parse the functional consequences of constitutive gene expression differences.

### Conclusion

Changing winters present many temperate ectotherms with a mosaic of new ecological circumstances. Warmer shorter winters might simultaneously indicate reductions in winter energetic requirements (M.J. Fitzpatrick et al., 2020), reductions in female fecundity (Benard, 2015), and potentially, novel immunological fitness landscapes. Adaptations to winters past might provide valuable information about how individuals will cope with future winters and climates. Our findings, which reveal large differences in constitutive immune gene expression profiles between populations from different origin winter climates, invite inquiry, especially from comparative and experimental standpoints. If the patterns we observed are representative of other wide-ranging terrestrial ectothermic vertebrates, and if they are the result of selection rather than processes of drift, then intraspecific genetic differentiation among populations may decline as winters warm.

## Acknowledgements

All egg masses were collected with permission from relevant state and federal agencies (Virginia Department of Game and Inland Fishes Permit # 070707, West Virginia Division of Natural Resources Permit # 2022.039, USFWS Research and Monitoring Special Use: Ohio River Islands Permit # 2202-02, New Hampshire Fish and Game Department (no permit #), Vermont Fish and Wildlife Department Permit # SR-2022-06). All animal care and experimental protocols were approved by the University of Connecticut Institutional Animal Care and Use Committee (IACUC) under protocols A19-045 and A22-037.

Funding for this research was provided by a NIAID Grant #1R01AI1236-5901A1 to Daniel Bolnick, the Trainor Fund and the Ralph M. Wetzel Endowment Fund for vertebrate research, both to the Department of Ecology and Evolutionary Biology and Connecticut State Museum of Natural History. Funding was also provided by the Center for Conservation and Biodiversity Fund for Research to the University of Connecticut Department of Ecology and Evolutionary Biology, the Society for Comparative and Integrative Biology Grant in Aid of Research, the Herpetologists’ League E.E. Williams Award, and the Society for the Study of Amphibians and Reptiles Dean Metter Award. The authors thank Dr. Sarah Knutie, Dr. Mark C. Urban, Dr. Christopher J. Sullivan, Dr. Eric Schultz, Dr. Max Zavell Jaime Neill, Frank M.S. Muzio, Leilani Duarte, Alexandra Pouliot, Hailey Baranowski, and Meagan Forbus, Adrienne Ell, and Nicole Dahrouge for their generous assistance with conceptualization, animal husbandry, field work, lab work, and figure generation. Finally, the authors thank Elaine Barr, the Burke family, and Jeff Coombes for their generous assistance locating and sampling egg masses in the field.

## Data and code availability

Raw and processed sequencing data are available from the Gene Expression Omnibus server from NCBI (accession # GSE266397). All R code used to produce the analyses and figures contained in the manuscript will be made available on **GitHub** and all metadata used in the analysis will be made available from **FigShare** upon acceptance of the manuscript.

