## Supplemental Materials for "Constitutive differences in immune gene expression and energy storage phenotypes co-vary with winter environment in wood frogs"

**Table S1.** Enriched biological process GO terms in up- and downregulated differentially expressed (DE) gene sets. Up to 10 GO terms with (BH-adjusted p≤0.05) are shown for each comparison.

| Comparison | Expression of genes with enrichment | GO Term | Description | Number DE genes in category | Number genes in category in background dataset | BH-adjusted P value |
|---|---|---|---|---|---|---|
| Harsh (-2°C): Mild vs. Harsh origin | Up | GO:0009410 | response to xenobiotic stimulus | 26 | 155 | < 0.001 |
| | | GO:0006805 | xenobiotic metabolic process | 23 | 147 | 0.005 |
| | | GO:0071466 | cellular response to xenobiotic stimulus | 23 | 147 | 0.005 |
| | Down | GO:0002252 | immune effector process | 45 | 238 | 0.025* |
| | | GO:1902105 | regulation of leukocyte differentiation | 30 | 139 | 0.040* |
| | | GO:0002819 | regulation of adaptive immune response | 20 | 71 | 0.025* |
| | | GO:0002577 | regulation of antigen processing and presentation | 7 | 10 | 0.025* |
| Harsh origin: Harsh (-2°C) vs. Mild (+2°C) | Up | GO:0034330 | cell junction organization | 8 | 207 | 0.028* |
| | | GO:0034329 | cell junction assembly | 7 | 172 | 0.035* |
| Harsh origin: Harsh (-2°C) vs. Pre-winter | Up | GO:0050789 | regulation of biological process | 586 | 5852 | 0.042* |
| | | GO:0019222 | regulation of metabolic process | 406 | 3794 | 0.005 |
| | | GO:0048519 | negative regulation of biological process | 301 | 2766 | 0.026* |
| | | GO:0009889 | regulation of biosynthetic process | 269 | 2411 | 0.013* |
| | | GO:0031326 | regulation of cellular biosynthetic process | 262 | 2384 | 0.038* |

| | | | | | | |
|---|---|---|---|---|---|---|
| | | GO:0010556 | regulation of macromolecule biosynthetic process | 259 | 2288 | 0.007 |
| | | GO:2000112 | regulation of cellular macromolecule biosynthetic process | 249 | 2227 | 0.021* |
| | | GO:0051252 | regulation of RNA metabolic process | 232 | 2084 | 0.043* |
| | | GO:0009792 | embryo development ending in birth or egg hatching | 216 | 1883 | 0.015* |
| | | GO:0031325 | positive regulation of cellular metabolic process | 187 | 1557 | 0.005 |
| | Down | GO:0044255 | cellular lipid metabolic process | 127 | 669 | 0.047* |
| | | GO:0044282 | small molecule catabolic process | 61 | 245 | 0.004 |
| | | GO:0006631 | fatty acid metabolic process | 58 | 248 | 0.029* |
| | | GO:0016042 | lipid catabolic process | 53 | 208 | 0.009 |
| | | GO:0019395 | fatty acid oxidation | 23 | 63 | 0.011* |
| | | GO:0034440 | lipid oxidation | 23 | 65 | 0.018* |
| | | GO:0006635 | fatty acid beta-oxidation | 16 | 40 | 0.043* |
| | | GO:0030049 | muscle filament sliding | 15 | 27 | 0.004 |
| | | GO:0033275 | actin-myosin filament sliding | 15 | 29 | 0.005 |
| Harsh origin: Mild (+2°C) vs. Pre-winter | Up | GO:0044238 | primary metabolic process | 671 | 5810 | 0.019* |
| | | GO:0044237 | cellular metabolic process | 668 | 5810 | 0.037* |
| | | GO:0034641 | cellular nitrogen compound metabolic process | 430 | 3525 | 0.013* |
| | | GO:1901360 | organic cyclic compound metabolic process | 428 | 3556 | 0.040* |
| | | GO:0006725 | cellular aromatic compound metabolic process | 417 | 3415 | 0.015* |
| | | GO:0046483 | heterocycle metabolic process | 416 | 3393 | 0.011* |

| | | | | | | |
|---|---|---|---|---|---|---|
| | | GO:0019219 | regulation of nucleobase-containing compound metabolic process | 316 | 2502 | 0.014* |
| | | GO:0009889 | regulation of biosynthetic process | 309 | 2438 | 0.014* |
| | | GO:0051252 | regulation of RNA metabolic process | 272 | 2105 | 0.012* |
| | | GO:0006355 | regulation of DNA-templated transcription | 255 | 2006 | 0.047* |
| Mild (+2°C): Mild vs. Harsh origin | Down | GO:0002252 | immune effector process | 30 | 248 | 0.013* |
| | | GO:0050867 | positive regulation of cell activation | 22 | 169 | 0.039* |
| | | GO:0002819 | regulation of adaptive immune response | 14 | 74 | 0.039* |
| Mild origin: Harsh (-2°C) vs. Mild (+2°C) | Down | GO:0048519 | negative regulation of biological process | 144 | 2095 | 0.045* |
| | | GO:0071310 | cellular response to organic substance | 74 | 913 | 0.045* |
| | | GO:0009887 | animal organ morphogenesis | 69 | 799 | 0.019* |
| | | GO:0071495 | cellular response to endogenous stimulus | 50 | 495 | 0.006 |
| | | GO:0009894 | regulation of catabolic process | 46 | 484 | 0.045* |
| | | GO:0006366 | transcription by RNA polymerase II | 40 | 390 | 0.034* |
| | | GO:0019433 | triglyceride catabolic process | 5 | 9 | 0.044* |
| Mild origin: Harsh (-2°C) vs. Pre-winter | Up | GO:0071704 | organic substance metabolic process | 639 | 5908 | 0.017* |
| | | GO:0044238 | primary metabolic process | 630 | 5810 | 0.016* |
| | | GO:0019222 | regulation of metabolic process | 431 | 3840 | 0.034* |
| | | GO:0060255 | regulation of macromolecule metabolic process | 357 | 3082 | 0.017* |
| | | GO:0010468 | regulation of gene expression | 297 | 2483 | 0.009 |

| | | | | | | |
|---|---|---|---|---|---|---|
| | | GO:0009889 | regulation of biosynthetic process | 288 | 2438 | 0.025* |
| | | GO:0031326 | regulation of cellular biosynthetic process | 285 | 2411 | 0.026* |
| | | GO:0010556 | regulation of macromolecule biosynthetic process | 273 | 2314 | 0.038* |
| | | GO:2000112 | regulation of cellular macromolecule biosynthetic process | 268 | 2252 | 0.028* |
| | | GO:0000003 | reproduction | 267 | 2241 | 0.026* |
| Mild origin: Mild (+2°C) vs. Pre-winter | Up | GO:0032501 | multicellular organismal process | 842 | 4944 | 0.021* |
| | | GO:0043170 | macromolecule metabolic process | 818 | 4793 | 0.023* |
| | | GO:0007275 | multicellular organism development | 768 | 4424 | 0.005 |
| | | GO:0044260 | cellular macromolecule metabolic process | 765 | 4392 | 0.004 |
| | | GO:0051716 | cellular response to stimulus | 621 | 3475 | 0.002 |
| | | GO:0048731 | system development | 619 | 3514 | 0.010* |
| | | GO:0048513 | animal organ development | 499 | 2767 | 0.009 |
| | | GO:0090304 | nucleic acid metabolic process | 475 | 2625 | 0.01 |
| | | GO:0006950 | response to stress | 434 | 2390 | 0.016* |
| | | GO:0009790 | embryo development | 417 | 2305 | 0.026* |
| | Down | GO:0048285 | organelle fission | 114 | 325 | 0.014* |
| | | GO:0000280 | nuclear division | 111 | 311 | 0.008 |
| | | GO:0044282 | small molecule catabolic process | 89 | 249 | 0.036* |
| | | GO:0016054 | organic acid catabolic process | 66 | 169 | 0.020* |
| | | GO:0046395 | carboxylic acid catabolic process | 66 | 169 | 0.020* |
| | | GO:0072329 | monocarboxylic acid catabolic process | 38 | 85 | 0.036* |

| | | | | | | |
|---|---|---|---|---|---|---|
| | | GO:0000236 | mitotic prometaphase | 34 | 62 | 0.002 |
| | | GO:0009062 | fatty acid catabolic process | 31 | 65 | 0.038* |
| Pre-winter: Mild vs. Harsh origin | Down | GO:0002376 | immune system process | 99 | 1221 | 0.033* |
| | | GO:0006955 | immune response | 58 | 605 | 0.023* |
| | | GO:0070663 | regulation of leukocyte proliferation | 18 | 109 | 0.023* |
| | | GO:0045619 | regulation of lymphocyte differentiation | 16 | 89 | 0.023* |
| | | GO:0045580 | regulation of T cell differentiation | 14 | 74 | 0.031* |
| | | GO:0042102 | positive regulation of T cell proliferation | 10 | 38 | 0.024* |
| | | GO:0002824 | positive regulation of adaptive immune response based on somatic recombination of immune receptors built from immunoglobulin superfamily domains | 9 | 33 | 0.032* |
| | | GO:0045577 | regulation of B cell differentiation | 7 | 19 | 0.030* |
| | | GO:0002579 | positive regulation of antigen processing and presentation | 5 | 9 | 0.032* |
| | | GO:0045591 | positive regulation of regulatory T cell differentiation | 4 | 5 | 0.030* |

**Table S2.** Genes expressed in an interaction between origin winter environment and artificial winter treatment.

| Predicted Gene | Description | Comparison | Pattern | Mean Expression | log2 FoldChange | SE | Stat | Padj |
|---|---|---|---|---|---|---|---|---|
| SLPI | XP_026264130.1 antileukoproteinase-like [Urocitellus parryii] | origin(Mild vs. Harsh): treatment (Freezing vs. Pre) | Negatively DE in freezing (-2°) versus pre artificial winter conditions in mild origin frogs but not harsh origin frogs | 24.2 | -3.9 | 0.9 | -4.2 | 0.1 |
| DDAH1 | XP_018417356.1 PREDICTED: N(G),N(G)-dimethylarginine dimethylaminohydrolase 1 isoform X1 [Nanorana parkeri] | origin(Mild vs. Harsh): treatment (Freezing vs. Pre) | Positively DE in freezing (-2°) versus pre artificial winter conditions in mild origin frogs but not harsh origin frogs | 9.0 | 2.0 | 0.5 | 4.1 | 0.1 |
| MOB3C | NP_001085692.1 MOB kinase activator 3C S homeolog [Xenopus laevis] | origin(Mild vs. Harsh): treatment (Freezing vs. Pre) | Negatively DE in freezing (-2°) versus pre artificial winter conditions in mild origin frogs but not harsh origin frogs | 1.6 | -3.4 | 0.8 | -4.3 | 0.1 |
| IFNGR2 | OCT91253.1 hypothetical protein XELAEV_18014304mg [Xenopus laevis] | origin(Mild vs. Harsh): treatment (Freezing vs. Pre) | Positively DE in freezing (-2°) versus pre artificial winter conditions in mild origin frogs but not harsh origin frogs | 5.9 | 2.0 | 0.4 | 4.6 | 0.0 |
| ITLN1 | sp\|Q5PPM0\|ITLN1_XENLA Intelectin-1 OS=Xenopus laevis OX=8355 GN=itln1 PE=1 SV=1 | origin(Mild vs. Harsh): treatment (Freezing vs. Pre); origin(Mild vs. Harsh): treatment | Positively DE in freezing (-2°) versus pre artificial winter conditions in mild origin frogs but not harsh origin frogs; Negatively DE in unfrozen (+2°C) versus freezing (-2°C) | 64.5 | 8.3; -5.6 | 1.4 | 6.0; -4.1 | <0.0001; 0.1 |

| | | | | | | | | |
|---|---|---|---|---|---|---|---|---|
| | | (Unfrozen vs. Freezing) | conditions in harsh origin but not mild origin frogs. | | | | | |
| HSPA8 | ACY69994.1 heat shock protein 70 [Pelophylax lessonae] | origin(Mild vs. Harsh): treatment (Freezing vs. Pre);origin(Mild vs. Harsh): treatment (Unfrozen vs. Pre) | Positively DE in both cold unfrozen (+2°C) and freezing (-2°C) conditions as compared to 'pre' in mild origin but not harsh origin frogs. | 9.6 | 5.9; 5.1 | 1.2 | 5.0; 4.4 | 0.003; 0.05 |
| MLL3 | XP_018408622.1 PREDICTED: histone-lysine N-methyltransferase 2C [Nanorana parkeri] | origin(Mild vs. Harsh): treatment (Unfrozen vs. Freezing) | Negatively DE in freezing (-2°) versus unfrozen (+2°C) artificial winter conditions in mild origin frogs but not harsh origin frogs | 14.2 | 1.1 | 0.3 | 4.0 | 0.1 |
| CDH2 | sp\|P30944\|CADH1_XENLA Cadherin-1 OS=Xenopus laevis OX=8355 GN=cdh1 PE=1 SV=2 | origin(Mild vs. Harsh): treatment (Unfrozen vs. Freezing) | Negatively DE in freezing (-2°) versus unfrozen (+2°C) artificial winter conditions in mild origin frogs but not harsh origin frogs | 47.7 | 0.8 | 0.2 | 4.0 | 0.1 |
| PHLDB3 | PIO31687.1 hypothetical protein AB205_0015920 [Lithobates catesbeianus] | origin(Mild vs. Harsh): treatment (Unfrozen vs. Freezing) | Negatively DE in freezing (-2°) versus unfrozen (+2°C) artificial winter conditions in mild origin frogs but not harsh origin frogs | 9.6 | 1.3 | 0.3 | 4.2 | 0.1 |

| | | | | | | | | |
|---|---|---|---|---|---|---|---|---|
| INF2 | sp\|Q0IHV1\|INF2_XENTR Inverted formin-2 OS=Xenopus tropicalis OX=8364 GN=inf2 PE=2 SV=1 | origin(Mild vs. Harsh): treatment (Unfrozen vs. Freezing) | Positively DE in freezing (-2°) versus unfrozen (+2°C) artificial winter conditions in harsh origin frogs but not mild origin frogs | 3.0 | 2.7 | 0.6 | 4.3 | 0.1 |
| NIPSNAP 1 | PIO16219.1 hypothetical protein AB205_0134080 [Lithobates catesbeianus] | origin(Mild vs. Harsh): treatment (Unfrozen vs. Pre) | Negatively DE in unfrozen (+2°C) versus 'pre' conditions in mild origin frogs but not harsh origin frogs | 3.8 | -1.9 | 0.4 | -4.2 | 0.1 |
| none identified | NP_001001229.1 stress-70 protein, mitochondrial [Xenopus tropicalis] | origin(Mild vs. Harsh): treatment (Unfrozen vs. Pre) | Positively DE in unfrozen (+2°C) versus 'pre' conditions in harsh origin frogs but not mild origin frogs | 40.4 | -1.1 | 0.2 | -4.4 | 0.1 |
| none identified | PIO30517.1 hypothetical protein AB205_0081970 [Lithobates catesbeianus] | origin(Mild vs. Harsh): treatment (Unfrozen vs. Pre) | Negatively DE in unfrozen (+2°C) versus 'pre' conditions in harsh origin frogs but not mild origin frogs | 0.7 | 5.1 | 1.1 | 4.5 | 0.1 |

**Table S3.** Differentially expressed (fdr-corrected P-value < 0.1) genes (up to 10 in each direction) for each comparison, with predicted gene symbol, log2fold change estimates.

| Comparison | Direction | Predicted gene | gene ID | log2Fold Change | lfc SE | FDR |
|---|---|---|---|---|---|---|
| Interaction: Origin X Treatment (Harsh vs. Pre) | Up | ITLN1 | MSTRG.29886 | 7.76 | 1.35 | < 0.001 |

| | | | | | | |
|---|---|---|---|---|---|---|
| | | HSPA8 | MSTRG.11304 | 5.16 | 1.89 | 0.003 |
| | | IFNGR2 | MSTRG.7069 | 1.06 | 0.75 | 0.020* |
| | | DDAH1 | MSTRG.34460 | 0.47 | 0.68 | 0.093 |
| | Down | MOB3C | MSTRG.34856 | -1 | 1.5 | 0.051 |
| | | SLPI | MSTRG.15799 | -0.82 | 1.55 | 0.073 |
| Interaction: Origin X Treatment (Mild vs. Harsh) | Up | PHLDB3 | MSTRG.40617 | 0.47 | 0.22 | 0.096 |
| | | CDH2 | MSTRG.37867 | 0.46 | 0.17 | 0.099 |
| | | MLL3 | MSTRG.28198 | 0.44 | 0.21 | 0.099 |
| | | INF2 | MSTRG.48505 | 0.28 | 0.24 | 0.096 |
| | Down | ITLN1 | MSTRG.29886 | -0.11 | 0.21 | 0.096 |
| Interaction: Origin X Treatment (Mild vs. Pre) | Up | HSPA8 | MSTRG.11304 | 0.1 | 0.18 | 0.051 |
| | Down | NIPSNAP1 | MSTRG.2105 | -0.3 | 0.27 | 0.08 |
| Harsh (-2°C): Mild vs. Harsh origin | Up | RPL14 | MSTRG.9631 | 5.5 | 1 | < 0.001 |
| | | HSDL1 | MSTRG.38439 | 4.83 | 0.45 | < 0.001 |
| | | RPL18A | MSTRG.23348 | 4.74 | 0.66 | < 0.001 |
| | | KIAA1704 | MSTRG.10051 | 4.22 | 0.79 | < 0.001 |
| | | PNLIP | MSTRG.42936 | 4.22 | 0.59 | < 0.001 |
| | | CHST9 | MSTRG.25828 | 4.11 | 0.7 | < 0.001 |
| | | NNMT | MSTRG.41766 | 3.74 | 0.44 | < 0.001 |
| | | OSBPL10 | MSTRG.27872 | 3.71 | 0.67 | < 0.001 |
| | | HSPA8 | MSTRG.11304 | 3.66 | 1.14 | < 0.001 |
| | | A2ML1 | MSTRG.43161 | 3.64 | 1.52 | < 0.001 |
| | Down | DUS1L | MSTRG.46557 | -2.92 | 1.13 | < 0.001 |
| | | SRP14 | MSTRG.49034 | -2.9 | 1.71 | 0.003 |
| | | IGHE | MSTRG.2378 | -2.63 | 0.53 | < 0.001 |
| | | CEBPB | MSTRG.44661 | -2.55 | 0.57 | < 0.001 |
| | | FBXL14 | MSTRG.21768 | -2.38 | 0.63 | < 0.001 |
| | | IL2RG | MSTRG.35079 | -2.27 | 1.14 | 0.002 |
| | | SPRY1 | MSTRG.4440 | -2.25 | 0.9 | < 0.001 |

| | | | | | | |
|---|---|---|---|---|---|---|
| | | TAF4B | MSTRG.44890 | -2.25 | 0.65 | < 0.001 |
| | | TCAM1 | MSTRG.20923 | -2.19 | 0.63 | < 0.001 |
| | | SULT6B1 | MSTRG.13087 | -2.15 | 1.12 | 0.003 |
| Harsh origin: Harsh (-2°C) vs. Mild (+2°C) | Up | BCAS1 | MSTRG.44603 | 4.02 | 0.3 | < 0.001 |
| | | ALAS1 | MSTRG.36676 | 0.85 | 0.47 | 0.007 |
| | | KRT4 | MSTRG.15099 | 0.8 | 0.3 | 0.006 |
| | | INF2 | MSTRG.48505 | 0.78 | 0.68 | 0.014* |
| | | TFAP2B | MSTRG.15492 | 0.7 | 0.23 | 0.006 |
| | | DST | MSTRG.14176 | 0.7 | 0.17 | 0.001 |
| | | TRPS1 | MSTRG.25920 | 0.68 | 0.33 | 0.021* |
| | | TFAP4 | MSTRG.31122 | 0.62 | 0.3 | 0.027* |
| | | HBB | MSTRG.30478 | 0.61 | 0.36 | 0.030* |
| | | DSP | MSTRG.26929 | 0.61 | 0.16 | 0.002 |
| | Down | GPNMB | MSTRG.28062 | -0.54 | 0.3 | 0.043* |
| | | CTSE | MSTRG.40879 | -0.53 | 0.73 | 0.032* |
| | | MASTL | MSTRG.28347 | -0.53 | 0.39 | 0.043* |
| | | GPR146 | MSTRG.30880 | -0.52 | 0.23 | 0.032* |
| | | VMP1 | MSTRG.9158 | -0.52 | 0.33 | 0.051 |
| | | SGMS1 | MSTRG.43415 | -0.52 | 0.22 | 0.028* |
| | | IL17D | MSTRG.46209 | -0.51 | 0.22 | 0.030* |
| | | MSMB | MSTRG.43629 | -0.51 | 0.29 | 0.054 |
| | | RAB32 | MSTRG.11176 | -0.5 | 0.41 | 0.048* |
| | | ARRDC3 | MSTRG.1202 | -0.5 | 0.26 | 0.05 |
| Harsh origin: Harsh (-2°C) vs. Pre-winter | Up | BCAS1 | MSTRG.44603 | 3.05 | 0.3 | < 0.001 |
| | | SFTPC | MSTRG.42048 | 2.3 | 0.45 | < 0.001 |
| | | ZFYVE20 | MSTRG.32599 | 2.01 | 0.31 | < 0.001 |
| | | D10JHU81E | MSTRG.35023 | 1.97 | 0.38 | < 0.001 |
| | | MORC3 | MSTRG.7018 | 1.91 | 0.41 | < 0.001 |
| | | PI3 | MSTRG.15825 | 1.78 | 0.58 | < 0.001 |

| | | | | | | |
|---|---|---|---|---|---|---|
| | | ISYNA1 | MSTRG.3570 | 1.69 | 0.42 | < 0.001 |
| | | WDR12 | MSTRG.30233 | 1.69 | 0.42 | < 0.001 |
| | | ARNTL | MSTRG.39691 | 1.68 | 0.59 | < 0.001 |
| | | HYAL2 | MSTRG.32474 | 1.67 | 0.66 | < 0.001 |
| | Down | CDC20 | MSTRG.34777 | -1.88 | 0.76 | < 0.001 |
| | | ABTB1 | MSTRG.32714 | -1.85 | 0.29 | < 0.001 |
| | | CCNB3 | MSTRG.35242 | -1.84 | 0.57 | < 0.001 |
| | | COMMD1 | MSTRG.12666 | -1.82 | 0.29 | < 0.001 |
| | | MUTYH | MSTRG.34953 | -1.79 | 0.71 | < 0.001 |
| | | H1F0 | MSTRG.33267 | -1.79 | 0.24 | < 0.001 |
| | | CASP9 | MSTRG.40320 | -1.78 | 0.51 | < 0.001 |
| | | DAK | MSTRG.39240 | -1.78 | 0.71 | < 0.001 |
| | | ZNF410 | MSTRG.48251 | -1.78 | 0.55 | < 0.001 |
| | | PAPSS2 | MSTRG.4310 | -1.77 | 0.52 | < 0.001 |
| Harsh origin: Mild (+2°C) vs. Pre-winter | Up | IRG1 | MSTRG.35005 | 2.73 | 0.82 | < 0.001 |
| | | HSPA13 | MSTRG.7211 | 2.64 | 0.66 | < 0.001 |
| | | D10JHU81E | MSTRG.35023 | 2.48 | 0.38 | < 0.001 |
| | | SLPI | MSTRG.15881 | 2.45 | 1.26 | < 0.001 |
| | | ZFYVE20 | MSTRG.32599 | 2.22 | 0.31 | < 0.001 |
| | | PI3 | MSTRG.15825 | 2.18 | 0.58 | < 0.001 |
| | | HS3ST5 | MSTRG.5268 | 2.17 | 0.41 | < 0.001 |
| | | RNF183 | MSTRG.37544 | 2.14 | 0.52 | < 0.001 |
| | | HYAL2 | MSTRG.32474 | 2.12 | 0.66 | < 0.001 |
| | | SFTPC | MSTRG.42048 | 2.12 | 0.47 | < 0.001 |
| | Down | TMEM42 | MSTRG.27718 | -2.13 | 0.84 | < 0.001 |
| | | RIPK3 | MSTRG.49276 | -2.1 | 0.7 | < 0.001 |
| | | MUTYH | MSTRG.34953 | -2.09 | 0.72 | < 0.001 |
| | | NKIRAS1 | MSTRG.27926 | -2.08 | 0.34 | < 0.001 |
| | | TOP2A | MSTRG.45389 | -2.08 | 0.64 | < 0.001 |

| | | | | | | |
|---|---|---|---|---|---|---|
| | | ZNF689 | MSTRG.17687 | -2.08 | 0.65 | < 0.001 |
| | | AKR1A1 | MSTRG.21659 | -2.08 | 0.43 | < 0.001 |
| | | LDHD | MSTRG.38275 | -2.07 | 0.72 | < 0.001 |
| | | JMJD6 | MSTRG.46061 | -2.06 | 0.72 | < 0.001 |
| | | SLX1A | MSTRG.31669 | -2.05 | 0.74 | < 0.001 |
| Mild (+2°C): Mild vs. Harsh origin | Up | RPL14 | MSTRG.9631 | 8.19 | 1.03 | < 0.001 |
| | | KIAA1704 | MSTRG.10051 | 4.64 | 0.82 | < 0.001 |
| | | HSDL1 | MSTRG.38439 | 4.4 | 0.46 | < 0.001 |
| | | PNLIP | MSTRG.42936 | 3.78 | 0.58 | < 0.001 |
| | | FGGY | MSTRG.34722 | 3.6 | 1.92 | 0.002 |
| | | NNMT | MSTRG.41766 | 3.55 | 0.45 | < 0.001 |
| | | RPL18A | MSTRG.23348 | 3.46 | 0.71 | < 0.001 |
| | | PTPN9 | MSTRG.24259 | 3.24 | 0.96 | < 0.001 |
| | | CHST9 | MSTRG.25828 | 3.14 | 0.79 | < 0.001 |
| | | OXLD1 | MSTRG.46739 | 3.01 | 1.36 | 0.002 |
| | Down | FCER1G | MSTRG.47721 | -2.83 | 0.57 | < 0.001 |
| | | MMP3 | MSTRG.11025 | -2.78 | 1.21 | 0.002 |
| | | ZMAT2 | MSTRG.22318 | -2.52 | 0.97 | 0.001 |
| | | C4A | MSTRG.37386 | -2.49 | 1.29 | 0.004 |
| | | TSPYL2 | MSTRG.35152 | -2.43 | 0.94 | 0.001 |
| | | OTUB1 | MSTRG.38533 | -2.37 | 0.35 | < 0.001 |
| | | VWA5A | MSTRG.5887 | -2.2 | 0.36 | < 0.001 |
| | | TAF6 | MSTRG.20643 | -2.1 | 1.27 | 0.007 |
| | | A2ML1 | MSTRG.42742 | -2.03 | 0.61 | < 0.001 |
| | | NUDC | MSTRG.8020 | -2.02 | 1.08 | 0.005 |
| Mild origin: Harsh (-2°C) vs. Mild (+2°C) | Up | BCAS1 | MSTRG.44603 | 2.47 | 0.27 | < 0.001 |
| | | KIAA0101 | MSTRG.24307 | 1.66 | 0.96 | 0.001 |
| | | RTP4 | MSTRG.14306 | 1.34 | 1.87 | 0.004 |
| | | KLF10 | MSTRG.25975 | 0.77 | 0.26 | 0.002 |

| | | | | | | |
|---|---|---|---|---|---|---|
| | | PTCD3 | MSTRG.5513 | 0.68 | 0.32 | 0.007 |
| | | FUNDC1 | MSTRG.7907 | 0.66 | 0.24 | 0.004 |
| | | TRX-2 | MSTRG.2959 | 0.65 | 0.16 | < 0.001 |
| | | MRPL19 | MSTRG.15393 | 0.57 | 0.16 | < 0.001 |
| | | TP53RK | MSTRG.44533 | 0.53 | 0.38 | 0.026* |
| | | PRMT5 | MSTRG.2390 | 0.53 | 0.23 | 0.013* |
| | Down | TSC22D1 | MSTRG.10069 | -0.74 | 0.18 | < 0.001 |
| | | SLC2A1 | MSTRG.39924 | -0.73 | 0.27 | 0.003 |
| | | ZNF148 | MSTRG.45129 | -0.72 | 0.41 | 0.007 |
| | | HLX | MSTRG.12427 | -0.72 | 0.41 | 0.007 |
| | | UBXN2A | MSTRG.15355 | -0.71 | 0.16 | < 0.001 |
| | | NIPA2 | MSTRG.5038 | -0.71 | 0.29 | 0.005 |
| | | USP53 | MSTRG.4381 | -0.71 | 0.22 | 0.002 |
| | | LRRN4CL | MSTRG.38554 | -0.7 | 0.48 | 0.01 |
| | | GRASP | MSTRG.10110 | -0.7 | 0.32 | 0.006 |
| | | ELTD1 | MSTRG.34511 | -0.68 | 0.52 | 0.011* |
| Mild origin: Harsh (-2°C) vs. Pre-winter | Up | HSPA8 | MSTRG.11304 | 6.04 | 0.73 | < 0.001 |
| | | HSPA8 | MSTRG.48979 | 4.98 | 0.85 | < 0.001 |
| | | ZFYVE20 | MSTRG.32599 | 3.43 | 0.41 | < 0.001 |
| | | BCAS1 | MSTRG.44603 | 2.94 | 0.3 | < 0.001 |
| | | HSPA8 | MSTRG.44325 | 2.86 | 0.98 | < 0.001 |
| | | CYP1B1 | MSTRG.24291 | 2.58 | 0.63 | < 0.001 |
| | | CH25H | MSTRG.43473 | 2.57 | 0.57 | < 0.001 |
| | | ODC1 | MSTRG.15236 | 2.27 | 0.34 | < 0.001 |
| | | SFTPC | MSTRG.42048 | 2.14 | 0.51 | < 0.001 |
| | | TSC22D3 | MSTRG.35332 | 2.13 | 0.21 | < 0.001 |
| | Down | ABTB1 | MSTRG.32714 | -1.97 | 0.32 | < 0.001 |
| | | ZFHX2 | MSTRG.2405 | -1.96 | 0.38 | < 0.001 |
| | | WNT5A | MSTRG.32966 | -1.95 | 0.35 | < 0.001 |

| | | | | | | |
|---|---|---|---|---|---|---|
| | | SMUG1 | MSTRG.10493 | -1.94 | 0.53 | < 0.001 |
| | | C11ORF57 | MSTRG.41331 | -1.92 | 0.36 | < 0.001 |
| | | FAM83D | MSTRG.46274 | -1.92 | 0.36 | < 0.001 |
| | | CASP9 | MSTRG.40320 | -1.92 | 0.6 | < 0.001 |
| | | MBOAT1 | MSTRG.15207 | -1.9 | 0.45 | < 0.001 |
| | | PLK1 | MSTRG.31155 | -1.89 | 0.38 | < 0.001 |
| | | CYSTM1 | MSTRG.22608 | -1.87 | 0.27 | < 0.001 |
| Mild origin: Mild (+2°C) vs. Pre-winter | Up | HSPA8 | MSTRG.11304 | 5.37 | 0.76 | < 0.001 |
| | | BHLHE40 | MSTRG.32275 | 3.79 | 0.5 | < 0.001 |
| | | HSPA8 | MSTRG.48979 | 3.63 | 0.98 | < 0.001 |
| | | ZFYVE20 | MSTRG.32599 | 3.58 | 0.39 | < 0.001 |
| | | HYAL2 | MSTRG.32474 | 3.45 | 0.69 | < 0.001 |
| | | ODC1 | MSTRG.15236 | 3.34 | 0.33 | < 0.001 |
| | | FOSL1 | MSTRG.38632 | 3.2 | 0.47 | < 0.001 |
| | | CYP1B1 | MSTRG.24291 | 3.08 | 0.54 | < 0.001 |
| | | PPIF | MSTRG.43889 | 2.65 | 0.43 | < 0.001 |
| | | HSPA13 | MSTRG.7211 | 2.64 | 0.57 | < 0.001 |
| | Down | C14ORF79 | MSTRG.48603 | -2.33 | 0.56 | < 0.001 |
| | | SMC2 | MSTRG.3055 | -2.3 | 0.65 | < 0.001 |
| | | C16ORF58 | MSTRG.31847 | -2.27 | 0.42 | < 0.001 |
| | | KLF15 | MSTRG.8434 | -2.27 | 0.59 | < 0.001 |
| | | ACSS3 | MSTRG.23246 | -2.25 | 0.31 | < 0.001 |
| | | FUT1 | MSTRG.41480 | -2.24 | 0.41 | < 0.001 |
| | | F3 | MSTRG.33846 | -2.23 | 0.26 | < 0.001 |
| | | C16ORF87 | MSTRG.38065 | -2.19 | 0.3 | < 0.001 |
| | | SETDB2 | MSTRG.10923 | -2.18 | 0.61 | < 0.001 |
| | | GGACT | MSTRG.9828 | -2.17 | 0.67 | < 0.001 |
| Pre-winter: Mild vs. Harsh origin | Up | A2ML1 | MSTRG.43161 | 5.75 | 1.14 | < 0.001 |
| | | RPL14 | MSTRG.9631 | 5.55 | 0.99 | < 0.001 |

| | | | | | |
|---|---|---|---|---|---|
| | RPL18A | MSTRG.23348 | 5.33 | 0.66 | < 0.001 |
| | KIAA1704 | MSTRG.10051 | 5.24 | 0.82 | < 0.001 |
| | PNLIP | MSTRG.42936 | 3.87 | 0.58 | < 0.001 |
| | NNMT | MSTRG.41766 | 3.42 | 0.45 | < 0.001 |
| | FGGY | MSTRG.34722 | 3.1 | 1.96 | 0.003 |
| | HSDL1 | MSTRG.38439 | 3.02 | 0.45 | < 0.001 |
| | OXLD1 | MSTRG.46739 | 3.02 | 1.33 | 0.001 |
| | SULT1C2 | MSTRG.31640 | 2.83 | 1.45 | 0.002 |
| Down | AMPD3 | MSTRG.39740 | -2.46 | 0.4 | < 0.001 |
| | FCER1G | MSTRG.47721 | -2.45 | 0.5 | < 0.001 |
| | GTPBP4 | MSTRG.28256 | -2.44 | 0.7 | < 0.001 |
| | A2ML1 | MSTRG.42742 | -2.35 | 0.54 | < 0.001 |
| | NCKAP1 | MSTRG.10466 | -2.27 | 1.14 | 0.003 |
| | TGFBI | MSTRG.22700 | -2.26 | 1.03 | 0.003 |
| | HBA | MSTRG.30476 | -2.08 | 0.48 | < 0.001 |
| | OTUB1 | MSTRG.38533 | -2.03 | 0.33 | < 0.001 |
| | FAM136A | MSTRG.21441 | -2.02 | 0.48 | < 0.001 |
| | TAF6 | MSTRG.20643 | -1.95 | 1.18 | 0.007 |

**Supplemental Methods**

*WGCNA*

We identified groups of genes (modules) whose expression changed in a manner correlated to the values of explanatory variables in our dataset. To do this, we performed weighted gene co-expression network analysis (WGCNA) using the *WGCNA* (v. 1.72-1) package in R (Zhang and Horvath, 2005). As input data, we used variance stabilized transformed data (generated with the 'vst' function in the *DESeq2* package) and generated signed networks. We used a soft thresholding power of 3, and a minimum module size of 100 for networks and merged modules with a dissimilarity threshold below 0.25 (Figure S1). We then used the the 'lmFit' and 'eBayes' functions from the R package *limma* (v.3.54.2) to model the expression of each module's eigengene in response to the same explanatory variables used in DESeq2 analysis (Ritchie et al., 2015). This analysis produced foldchange estimates and (Benjamini-Hochberg adjusted) significance estimates for each module eigengene. We used a threshold of adjusted p $<0.01$, and $|\text{Pearson's } R| \geq 0.7$ to determine whether the correlation between module-wide expression and at least one explanatory variable in the dataset investigated modules was significant.

We used the 'goseq' (v 1.5.0) package (Young et al., 2010) in R to identify enriched gene ontology (GO) terms in lists of differentially expressed genes from DESeq2 analysis, and in lists of genes belonging to modules identified by WGCNA as having the most significant correlations (as defined previously) with explanatory variables. Importantly, differentially downregulated gene sets can be enriched for GO terms; in these cases GO terms are significantly overrepresented among genes with significant negative log2fold changes. All enrichment tests included a probability-weighting function to correct for the biasing effect of transcript length on the probability that a gene is differentially expressed. The probability weighting function was constructed using the *pwf* function and input lists of differentially expressed genes and their corresponding transcript lengths. Enriched GO terms with a Benjamini-Hochberg adjusted p-value <0.05 were considered significant. For visualizations and to focus our analysis on changes in immune processes, we kept all terms descended from the immune system process (GO:0002376), innate immune response (GO:0045087), and the adaptive immune response (GO:0002250) parent GO terms. We present data on how biological processes in these parent categories vary in our study, and present more general results on significantly enriched biological processes (not just immune processes) in the supplementary information materials.

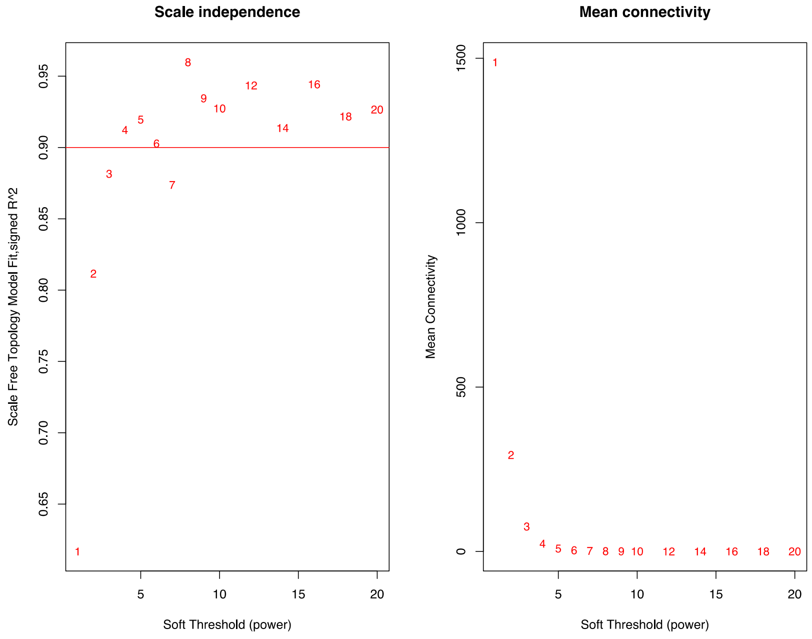


**Figure S1**. WGCNA thresholding and connectivity.

### Supplemental Results

*WGCNA*

Weighted gene co-expression network analysis identified 13 gene modules (Figure S2). Of these, six were significantly correlated with at least one explanatory variable in our dataset. Expression of module genes was generally correlated with origin environment, artificial winter treatment, or their interaction (Figure S3, Table S3).

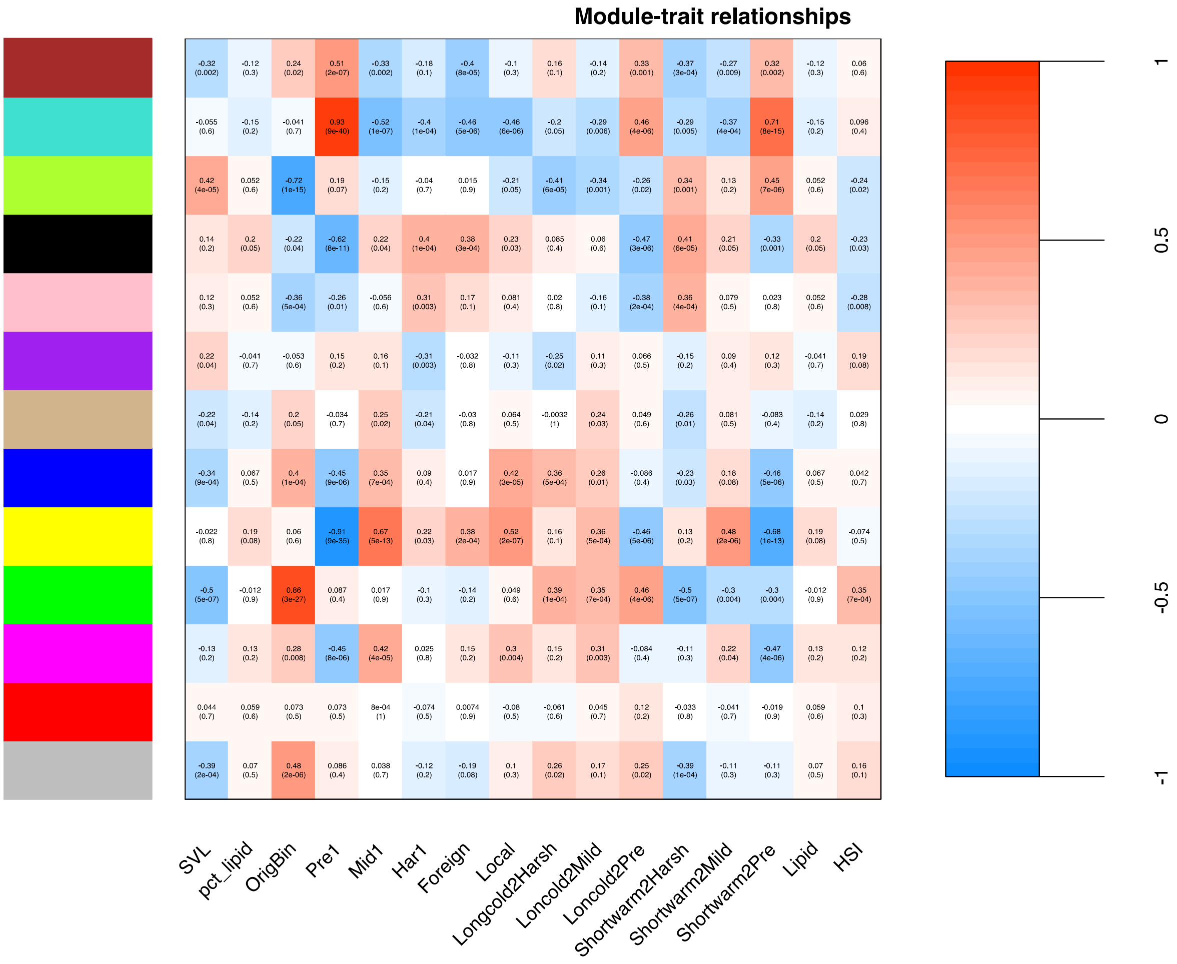


**Figure S2.** Thirteen gene modules were identified from wood frog ventral skin, of which 7 were significantly correlated (|*Pearson's* R| > 0.7)with variables in our dataset.

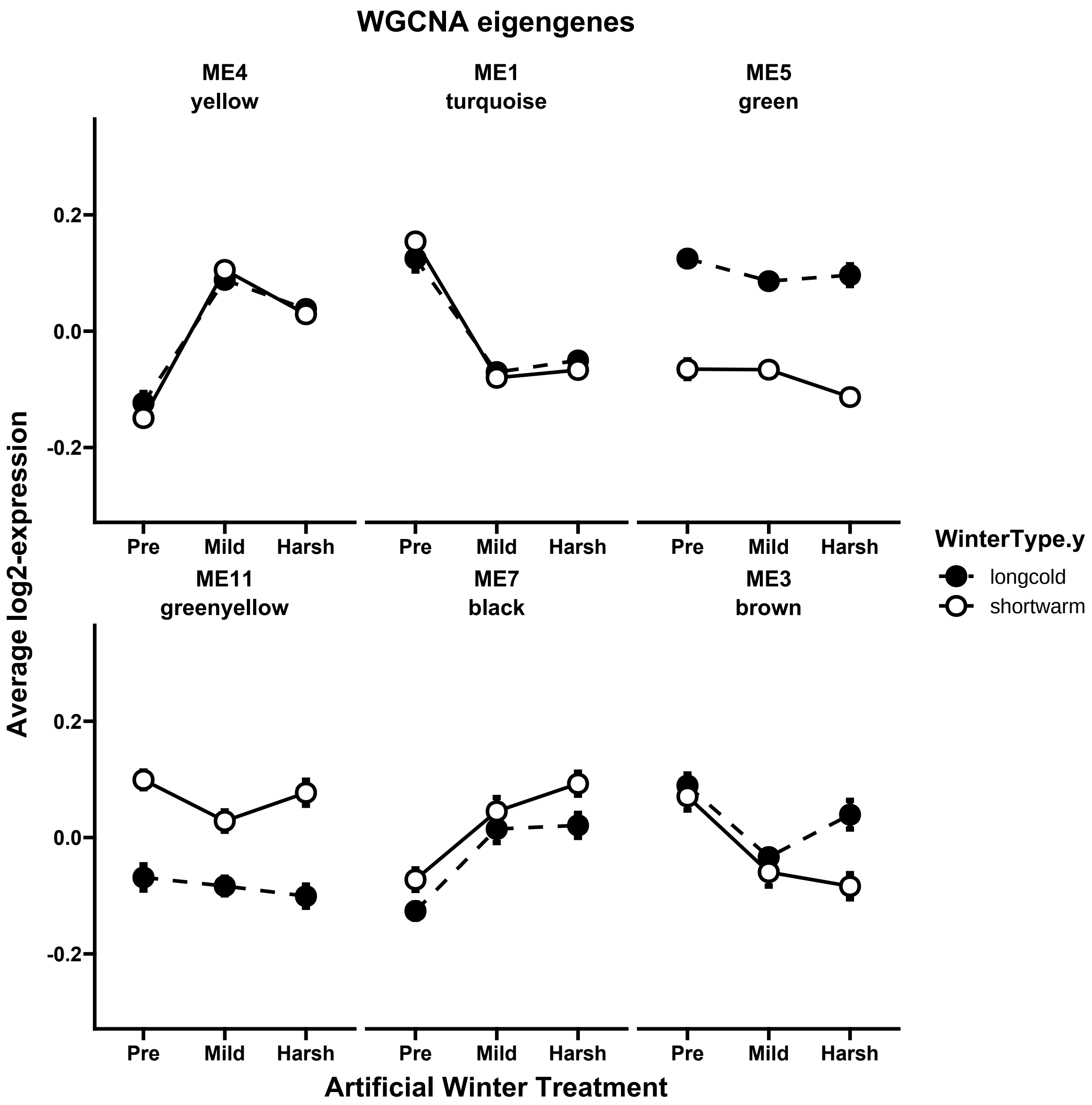


**Figure S3.** Expression of module eigengenes identified with WGCNA. Modules varied in expression in response to treatment (yellow, turquoise, black), frog origin environment (winter type, green, greenyellow) and with the interaction between treatment and frog origin (brown).

**Table S3.** Effect sizes and adjusted P-values for module eigengenes correlated with origin environment, treatment, and their interaction. Parameter estimates for explanatory variables and p-values for expression of module eigengenes with explanatory variables are displayed.

| | | | *Limma lmFit() estimated average log2 expression effect size* | | | | | | | |
|---|---|---|---|---|---|---|---|---|---|---|
| Module number | Module size | Module color | Mild origin | Harsh (-2°C) Treatment | Mild (+2°C) Treatment | Interaction: Origin X Harsh v Pre trt. | Interaction: Origin X Mild v Pre trt. | Ave Expr | F | B-H adjusted p-value |
| 4 | 850 | yellow | -0.026 | 0.162 | 0.211 | 0.017 | 0.043 | 8.73e-17 | 122.60 | < 0.001 |
| 1 | 2423 | turquoise | 0.029 | -0.175 | -0.195 | -0.046 | -0.039 | 2.08e-16 | 106.74 | < 0.001 |
| 5 | 751 | green | -0.19 | -0.028 | -0.039 | -0.019 | 0.038 | 1.92e-16 | 54.88 | < 0.001 |
| 11 | 140 | greenyellow | 0.168 | -0.032 | -0.015 | 0.01 | -0.056 | 3.69e-17 | 22.53 | < 0.001 |
| 7 | 306 | black | 0.054 | 0.147 | 0.141 | 0.018 | -0.023 | 8.71e-17 | 14.57 | < 0.001 |
| 2 | 882 | blue | -0.078 | 0.111 | 0.088 | -0.063 | 0.053 | 8.43e-17 | 12.29 | < 0.001 |
| 3 | 863 | brown | -0.019 | -0.05 | -0.123 | -0.104 | -0.007 | 1.33e-16 | 10.95 | < 0.001 |
| 9 | 262 | magenta | -0.079 | 0.06 | 0.098 | 0.016 | 0.052 | 6.81e-17 | 8.59 | < 0.001 |
| 0 | 4436 | grey | -0.091 | -0.004 | -0.024 | -0.061 | 0.024 | 2.97e-16 | 6.72 | < 0.001 |
| 8 | 298 | pink | 0.107 | 0.107 | 0.063 | -0.03 | -0.051 | 9.11e-17 | 6.13 | < 0.001 |
| 12 | 131 | tan | -0.031 | -0.014 | 0.045 | -0.027 | -0.009 | 5.61e-17 | 2.27 | 0.064 |
| 10 | 180 | purple | 0.008 | -0.078 | 0.01 | 0.019 | -0.016 | 1.64e-16 | 1.97 | 0.099 |
| 6 | 635 | red | -0.037 | -0.048 | -0.022 | 0.045 | 0.017 | 5.98e-17 | 0.36 | 0.873 |
